# Surface Patterned Omniphobic Tiles (SPOTs): a versatile platform for scalable liquid handling

**DOI:** 10.1101/2024.01.17.575712

**Authors:** Samira Shiri, Mohsin J Qazi, Shenghao Tan, Jon Albo, Arnold Chen, Rena Fukuda, Mika S Jain, Nkazi Nchinda, Mark Menesses, Ghada Ahmed, Arynn O Gallegos, Mahesh K Gangishetty, Daniel N Congreve, Nate J Cira

## Abstract

Manipulating liquids is a ubiquitous need for experiments across numerous scientific disciplines. To overcome limitations of current methods, we introduce Surface Patterned Omniphobic Tiles (SPOTs). This platform combines geometry and surface engineering, building on discontinuous wetting approaches to leverage capillarity for metering liquids. The SPOTs platform allows manipulation of hundreds to thousands of independent experiments without expensive equipment or large consumable costs. These devices can handle a wide range of liquid types and volumes (<10 nanoliters to >10 microliters) with better precision than pipetting. The platform is inexpensive and easy to fabricate, fast and intuitive to use, and cross-compatible with existing microwell plate layouts. We demonstrate how these capabilities facilitate diverse experiments including testing antibiotic combinations for synergy and antagonism, material screening of perovskites, and genotyping microbial isolates. We anticipate SPOTs will enable users from disparate domains to quickly and easily run a wide range of high-throughput experiments.

---

Experiments involving precisely manipulated liquid volumes play a central role in scientific discovery across numerous domains. The most widely adopted liquid handling approaches are manual and robotic pipetting which are intuitive and versatile but can be challenging to scale due to consumable costs, large volume requirements, and low throughput. Alternative liquid handling platforms have been developed, addressing the limitations of pipetting through diverse approaches including open^1^ and closed^2^ channels, emulsions^3–5^, on-chip valves^6,7^, and inkjet^8,9^ and acoustofluidic^10^ droplet ejection. These devices have transformed specific experimental tasks such as single-cell transcriptomics^9^, point-of-care immunoassays^11^, cell counting and sorting^12^, and the manipulation of large chemical compound libraries^13,14^. However, despite their sizable impacts, each existing liquid handling approach has a unique profile of advantages and disadvantages, and researchers often face substantial challenges when adapting them to new experimental workflows^15^.

A need exists for a platform that has both the accessibility and rapid adaptability of pipette-based approaches and the precise small-volume handling capabilities and high throughput of alternative approaches. Such a platform would facilitate more routine application of advanced liquid handling to diverse experimental workflows, which is especially important for the study of complex systems^16,17^ and for generating large datasets to support rapidly evolving data-driven approaches to prediction and discovery^18,19^.

Here we introduce Surface Patterned Omniphobic Tiles (SPOTs), which advance discontinuous dewetting approaches^20,21^ and microdroplet arrays^22^ by leveraging an omniphobic coating^23^ that repels a wide range of liquids, custom loading devices with carefully constructed geometries, and simple methods of fabrication and use. The resulting platform achieves precise and scalable liquid handling while remaining accessible and easily adaptable. We first describe the SPOTs platform – its fabrication, use, and liquid handling capabilities – then we demonstrate its advantages for three distinct experimental workflows.

## The SPOTs platform

The SPOTs platform has two main components: 1) SPOTs plates that selectively repel and attract liquids in predefined patterns, and 2) loading devices that retain and controllably deposit liquids by sliding across the SPOTs plates (Fig. 1A). To fabricate SPOTs, we developed an accessible workflow that does not require photolithographic masks, specialized equipment, or synthesis of custom molecules. We first sprayed coated glass substrates with an omniphobic^23^ coating made of fumed silica particles functionalized with fluorinated silane^24–26^. We then selectively removed the coating by ablation with a consumer-grade laser cutter to create-philic areas that meter precise liquid volumes. We fabricated the loading devices by milling them from plastic then applying the same omniphobic coating (fabrication details in Supplementary information section S1.1). We created loaders of two different types: “slot loaders” which load the same liquid across all the –philic regions of the SPOTs plate (Fig. 1A, S2a) and “row/column loaders” which can load different liquids in each row/column of the plate (Fig. 1D, S2b). The omniphobic coating allows these devices to repel liquids of high and low surface tensions (Fig. S1), extending the applicability of the platform to a wider variety of experiments.

**Fig. 1.**
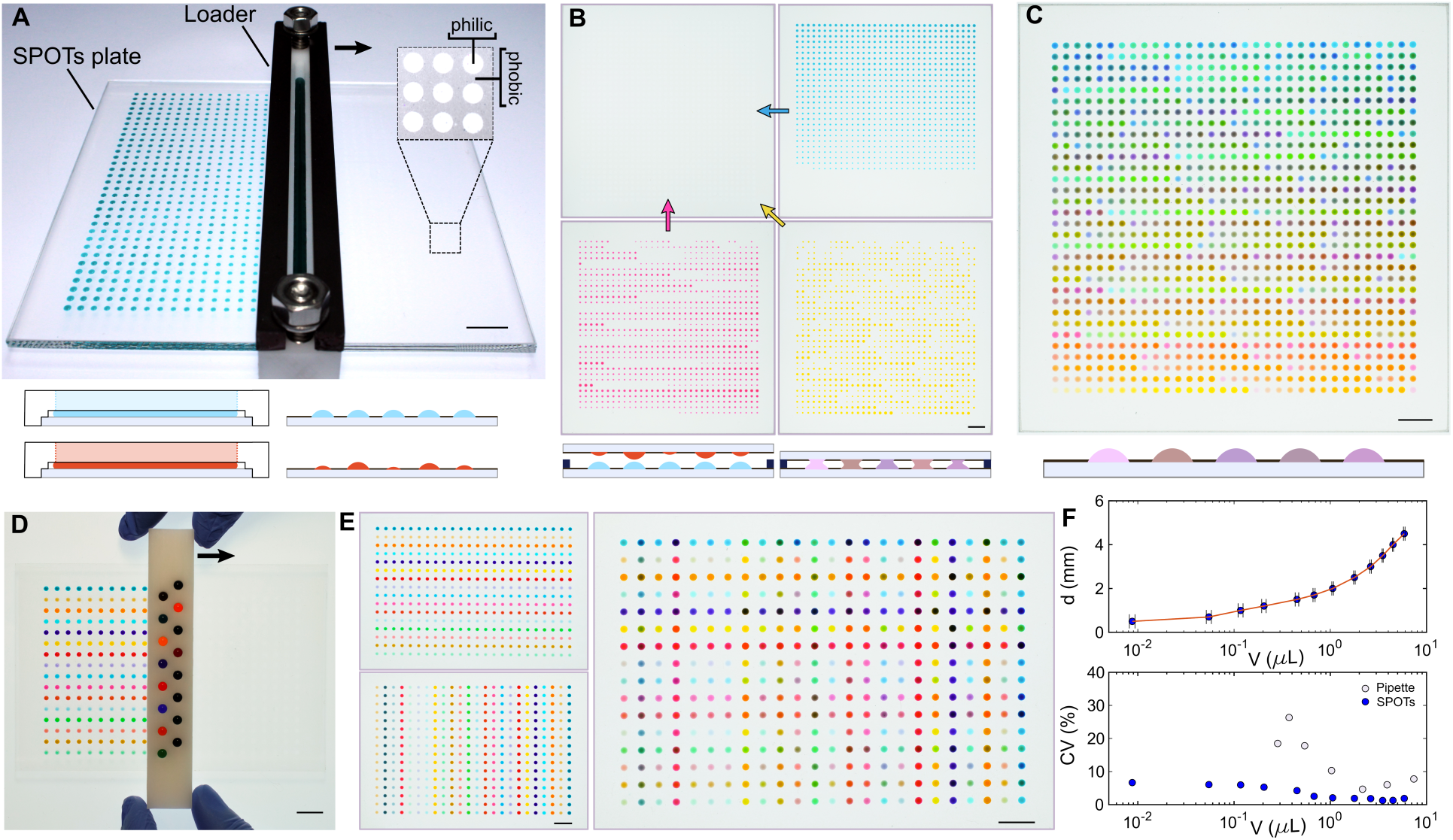
Liquid handling using the SPOTs platform. **a**, To load reagents, a loading device is filled and slid over the SPOTs plate. Each –philic circular area attracts liquid while the omniphobic surroundings repel it. Liquid volume is metered precisely by circle size. The arrow shows the sliding direction of the loader. **b,** Multiple plates can be loaded with different liquids which can be combined with sequential sandwiching to create target multi-liquid mixtures (curved arrows indicate flipping and sandwiching of plates). **c,** The workflow from (**a-c**) allows over 1,000 different 4-component liquid mixtures in sub-microliter volumes with four pipette tips in less than fifteen minutes (See also Movie S1). **d,** Modified loaders can deposit liquids on sub-sections of a plate (arrow shows loader sliding direction). **e,** This enables a different liquid to be loaded in each row and each column, which can be mixed to give pairwise combinations (See also Movie S2). **f,** Liquid handling is possible over three orders of magnitude in liquid volume (<10 nL to >10 μL). Compared to pipetting, SPOTs can handle a wider range of volumes and smaller volumes with less error. Error bars indicate one standard deviation. Coefficient of variation (CV) computed from n = 12 pipetted and n = 8 SPOTs volumes with the same device, see also Fig. S3. (Scale bars: 1 cm)

The SPOTs platform is capable of handling over three orders of magnitude of liquid volumes from 9 nL to 13 μL (Fig. 1F), with better precision than pipetting (Fig. 1F, S3). The deposited volume is set by the size of each patterned –philic area and the distance between the bottom of the loader and the top of the SPOTs plate. For high precision metering, loaders can be planarized with metal bars (Fig. 1A, Supplementary information section S1.3). The loaded volume is nearly invariant to differences in viscosity (Fig. S4A) and reasonable loading speeds (Fig. S5), while surface tension has a weak effect on the loaded volume (Fig. S4B).

Beyond metering single liquids, more complex, multistep liquid handling is easy with the SPOTs platform. Liquid reagents from different SPOTs plates can be combined by sandwiching plates together across a defined gap, causing droplets on opposing plates to merge and mix diffusively in the vertical, but not lateral, direction^20,27,28^ (Fig. 1B). This mixing can be sped up by bringing the plates closer and farther apart, through a Taylor dispersion-like mechanism (Movie S1). Higher order mixtures can be created by starting with a plate of uniformly arrayed liquid (a “receiver plate”) then sequentially mixing in additional liquids through sandwiching while taking into account the dilution that occurs from each subsequent addition. Movie S1 and Fig. 1, B and C demonstrate this process by creating 1,056 unique four-liquid mixtures in less than 15 minutes using only four pipette tips. It is also straightforward to create pairwise mixtures from a larger number of components using two plates and the row/column loaders (Fig. 1E). The number and layout of these liquid mixtures is customizable and can be designed for compatibility with existing equipment and footprints, such as microwell plates (Fig. 1E). To test how these capabilities generalize and enable experiments in different domains, we next executed three disparate integrated application workflows.

## Live cell growth and inhibition assays

Treatment of infections with combinations of antibiotics, rather than single agents, offers a strategy to increase antimicrobial efficacy and decrease the emergence of antimicrobial resistance^29,30^. However, evaluating the efficacy of drug combinations requires running a large number of inhibition assays involving different drugs, doses, and replicates, motivating the application of various high-throughput liquid handling approaches to this challenge^31–34^. Here, we demonstrate the utility of the SPOTs platform for performing high-throughput screening of pairwise antibiotic combinations to inhibit the growth of *Escherichia coli* W3110.

We evaluated pairwise interactions of five antibiotics: amoxicillin (AMX), polymyxin B (PMB), doxycycline (DOX), clarithromycin (CLA), and trimethoprim (TMP) (Supplementary Table I). Using a slot loader, media was loaded onto a SPOTs receiver plate with a uniform array of 2 mm diameter –philic areas. Again, using a slot loader, each antibiotic was sequentially loaded on a separate plate with a defined pattern then sandwiched and mixed into media on the receiver plate (Fig. 2A). *E. coli* W3110 was loaded on a separate plate, sandwiched with the receiver plate containing media and antibiotic combinations, and sealed with paraffin wax to prevent evaporation. This workflow consumed only seven pipette tips and resulted in 600 separate experiments on a single device. These included 480 pairwise drug combinations (four concentrations of five drugs with three replicates), 90 single drug experiments (various concentrations with three replicates, Fig. S6), 10 positive control experiments, and 20 negative control experiments. The sealed assembly was incubated at 35 °C on an automated microscope that measured the scattered light intensity with dark-field imaging every 30 minutes for 21 hours, to quantify cell growth (Fig. 2B, Detailed methods in Supplementary information section S2.1).

**Fig. 2.**
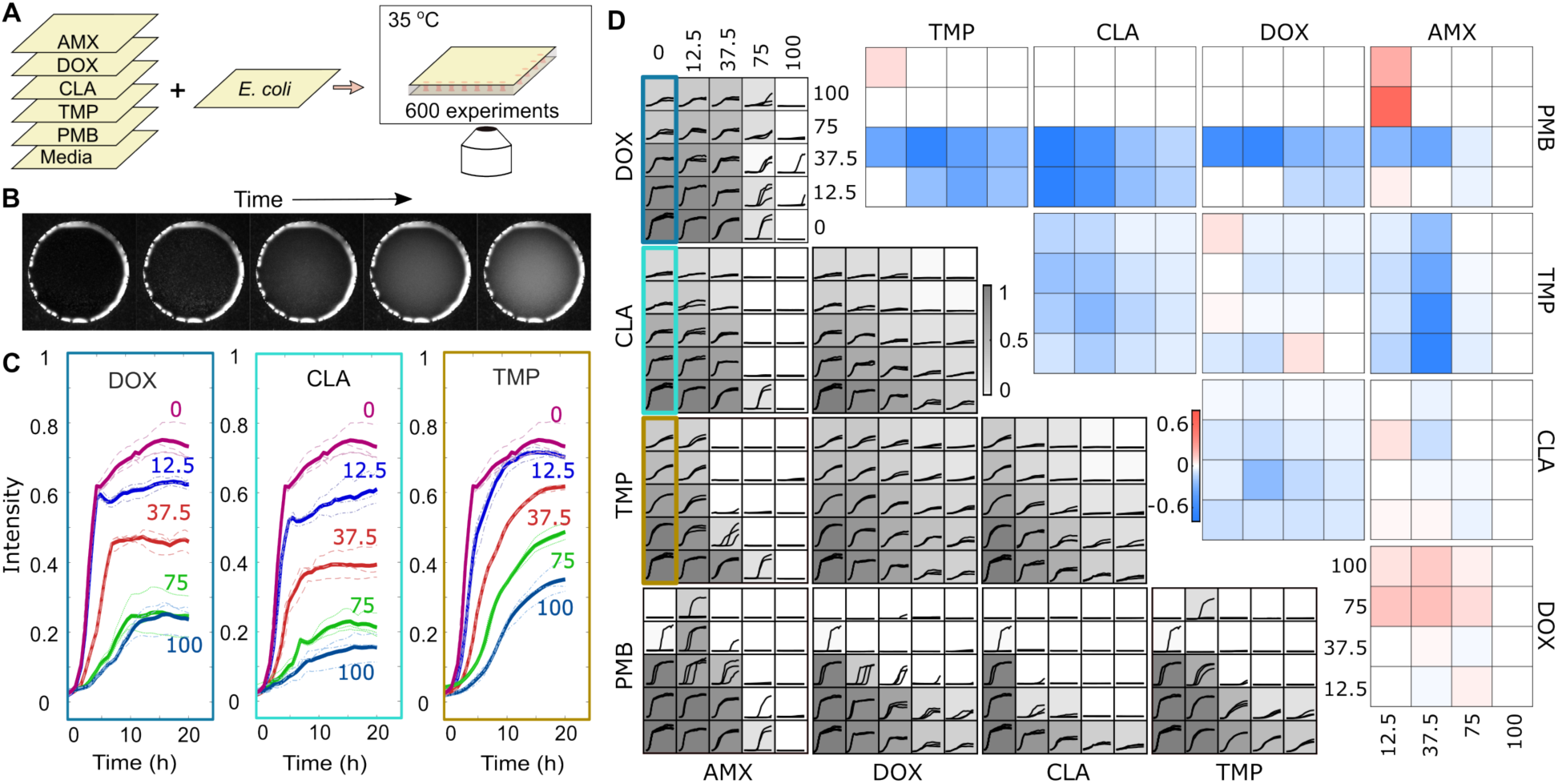
Screening antibiotic combinations using SPOTs. **a**, To run 600 experiments simultaneously, a receiver plate was loaded with media, then each antibiotic was loaded on a separate plate and sequentially added to the receiver plate by sandwiching. The resulting plate with antibiotic mixtures was sandwiched with a plate of *E. coli* cells, sealed with wax and incubated on the microscope stage. **b,** Time lapse dark-field imaging was used to image cell growth through an increase in turbidity over time within each 2 mm diameter –philic area. **c,** 24-hour growth curves of *E. Coli* in response to different antibiotic concentrations (percent maximum dose) for DOX, CLA, and TMP. **d,** Results of combinatorial checkerboard assays run on SPOTs. Left: Triplicate growth curves of different antibiotic combinations and doses (percent maximum dose); background grayscale indicates growth at 10.5 hours relative to no-antibiotic control. Right: Bliss scores indicating synergy (blue) and antagonism (red) at 10.5 hours, with other time points showing similar trends (Fig. S7).

Due to the parallel nature of SPOTs manipulations, a separate loading manipulation is required for each different antibiotic, but different doses, replicates, and pairwise combinations are obtained with little-to-no additional experimenter time or effort. For single antibiotics, we found that the growth of *E. Coli W3110* was inhibited in a dose-dependent manner (Fig. 2C). We also obtained a “checkerboard” of pairwise checkerboard assays (Fig. 2D, left), which we used to evaluate whether each drug interaction was antagonistic or synergistic, with the Bliss independence model^35^ shown in Fig. 2D (right). We found synergistic interactions between TMP/CLA, CLA/DOX, TMP/AMX, PMB/TMP, PMB/CLA, and PMB/DOX, while AMX/DOX was antagonistic. Several of these interactions have been previously observed^31,36^. Replicate experiments for some drugs were more consistent than others. Notably, PMB replicates were less consistent, aligning with previous observations for this drug^37,38^ and highlighting the importance of including replicates for robust inferences.

This application demonstrates biocompatibility, long-term incubation and imaging capabilities, and high-throughput small molecule screening with minute reagent quantities, establishing the SPOTs platform as a promising tool for numerous experiments involving live cell growth and inhibition.

## Combinatorial material screening

Lead-halide perovskites have emerged as a promising class of materials in applications such as cheap, efficient photovoltaic cells^39^ and light emitting diodes (LEDs)^40^. Perovskites have cubic or tetragonal crystal structures (Fig. 3A) and the general formula ABX_3_, where A and B are cations and X is an anion. Varying ionic composition can substantially alter material properties, such as changing the bandgap^41^ and improving stability^42,43^. The full combinatorial composition space is vast, motivating high-throughput efforts to connect compositions with properties^44–46^. Here we use SPOTs to explore this multidimensional composition space.

**Fig. 3.**
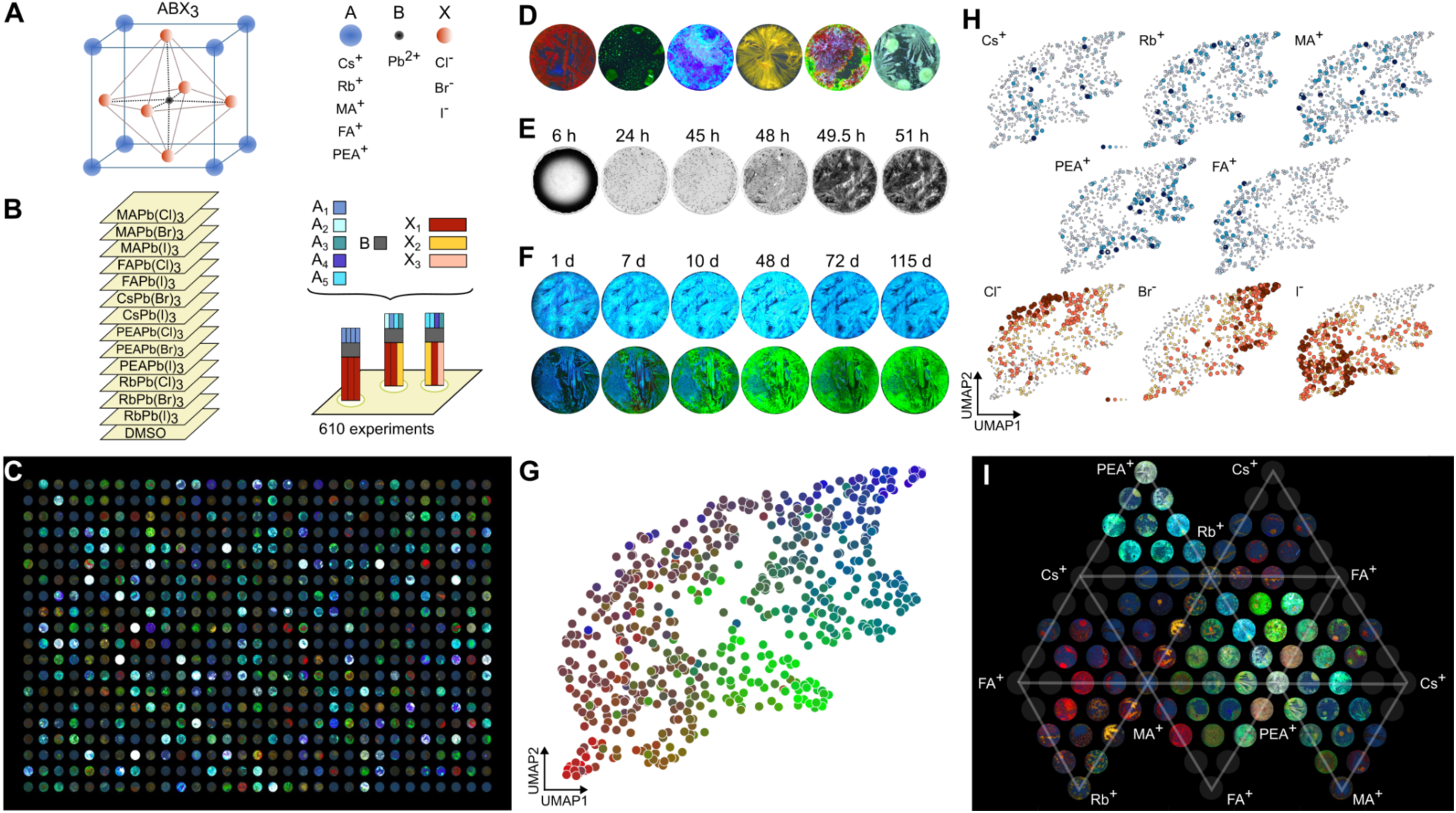
Combinatorial material creation using SPOTs. **a**, Perovskite crystal structure and ion substitutions explored. **b,** Experimental workflow schematic showing 13 precursor mixtures loaded onto different plates, and sequentially sandwiched onto a receiver plate on the right. Each spot can have up to four A site cations and 3 X site anions as indicated by different color of the bars. **c,** Stitched images of RGB emission of all materials on the SPOTs plate (374 nm excitation). **d,** Higher resolution images of materials with various colors, emissions, and morphologies. **e,** Time course images of a material forming during solvent evaporation (hours post-mixing). **f,** Aging in ambient conditions revealed consistent (top) and changing (bottom) materials. (Days post-formation; note: Minor changes due to fluctuations in excitation intensity.) **g,** UMAP projection showing a 2D view of the high dimensional material landscape. Each point represents a material, colored by its average intensity-normalized RGB value; images that appear more similar to each other are closer in space (Fig. S18). **h,** Mole fractions of each ion overlaid on the UMAP from (G). **i,** Ternary diagram of cation compositions for materials with equimolar anion ratios (Cl^-^, Br^-^, I^-^ 1:1:1). Gray circles indicate compositions that are missing due to solubility limitations.

We selected five A site cations (methylammonium (MA^+^), rubidium (Rb^+^), cesium (Cs^+^), formamidinium (FA^+^), and phenethylammonium (PEA^+^)), one B site cation (lead (Pb^2+^)), and three X site anions (chloride (Cl^-^), bromide (Br^-^) and iodide (I^-^)). To make different compositions, we dissolved precursors AX and BX_2_ in DMSO and then combined them to make 15 ABX_3_ stock solutions (5 A site cations with 3 X site anions). Two solutions (CsPbCl_3_ and FAPbBr_3_) were removed from the experiment due to poor solubility. We loaded a receiver plate with DMSO then loaded and sequentially sandwiched the 13 stock solutions onto the receiver plate (Fig. 3B). This created compositions with up to four A site cations (in increments of 25%) and up to three X site anions (in increments of 33%), resulting in 540 different compositions, 35 of which were created in triplicate, totaling to 610 mixtures. We then allowed the solvent to evaporate under ambient conditions (Fig. 3E) and imaged emission from the resulting materials using an RGB camera and UV excitation (374 nm) (Detailed methods in Supplementary information section S3.1). A diverse array of materials formed across the plate (Fig. 3C) including materials with homogenous emission of various colors, heterogenous emission, precursor precipitation, and a wide range of morphologies (Fig. 3D). Replicated compositions, distributed randomly across the plate, formed materials with similar appearance (Fig. S17). Additionally, select compositions recreated with pipettes resulted in materials with similar appearance to those created using SPOTs (Fig. S14). We note that some target compositions are expected to give materials with heterogeneous color or precursor precipitation due to limited uptake (e.g. high rubidium concentrations^47,48^), and for other compositions varying the solvent and crystallization conditions can yield more homogenous materials^49,50^.

To visualize this complex dataset, we extracted and vectorized 3D histograms of RGB color information from each image, then compressed this high dimensional information into 2D using Uniform Manifold Approximation and Projection (UMAP) (Fig. 3G). We overlaid the composition on this visualization (Fig. 3H), to reveal general trends. For example, compositions rich in PEA^+^ tended to yield blue and green crystals, as expected from the creation of layered materials leading to quantum confinement^51^. Br^-^ was associated with blue and green crystals, and I^-^ was associated with red crystals, agreeing with the bandgap changes associated with adjusting the halide species^52–54^. In addition to acquiring data immediately after material formation, we also acquired images of materials during solvent evaporation (Fig. 3E), and as materials aged in the months following solvent evaporation (Fig. 3F). During aging in ambient conditions, we observed a range of outcomes, including rapidly declining emission, consistent emission (Fig. 3F), and color change over time (Fig. 3F).

In addition to trends from holistic analysis of the entire dataset (Fig. 3, G and H), we also considered specific compositions with properties of particular interest. We curated a list of compositions that yielded materials with homogenous red, blue, or green emission (Fig S12, S13). Within this list, we found that the blue-emitting materials almost always contained PEA^+^ and always contained Br^-^, consistent with broad trends from figure 3H and quantum confinement resulting from PEA^+^ incorporation. Green-emitting materials had a wide variety of compositions, and all the homogenous red-emitting materials contained at least some FA^+^ and had both Br^-^ and I^-^. Further analysis of ternary diagrams revealed the impact of changes within reduced regions of the parameter space, such as varied cationic composition (Fig. 3I, S8-S11). From this analysis, we identified compositions with Cs^+^, MA^+^, and FA^+^ cations and equimolar anion ratios (Cl^-^, Br^-^, I^-^ 1:1:1) that produced materials with bright red emission, an important property for lighting applications. From these starting compositions, we performed a higher resolution sweep, and took photoluminescence measurements that confirmed the trends from the original experiment and augmented our understanding of these materials (Fig. S15, S16). We found peak emission wavelengths of 670 – 745 nm and high intensity emission for compositions with MA^+^, Cs^+^, and at least 20% FA^+^ (Fig S15, S16). Lead perovskite materials containing lower order subsets^55–57^ of these six ions are well appreciated, but such high complexity mixtures are less explored due to the mismatch between throughput of conventional methods and the number of combinatorial possibilities.

This application demonstrates how SPOTs can be used to generate combinatorial materials within high-dimensional composition spaces. The resulting rich data can provide insight into general trends as well as screen for materials with specific properties of interest. Such data can inform further SPOTs-coupled experiments that investigate select portions of the composition space at higher resolution or involve additional measurements such as photoluminescence quantum yield, emissive wavelength, and color purity.

## Genotyping microbial isolates

The 16S ribosomal RNA (rRNA) gene is commonly sequenced to identify bacteria and archaea^58^. The sequence of this region is useful for taxonomic identification^58,59^ and can even enable prediction of an organism’s metabolic capabilities^60^. Thus, the 16S rRNA gene is frequently sequenced in applications like microbial organism discovery^61,62^ and clinical isolate identification^63–65^. However, preparing microbial samples for sequencing involves numerous, often repetitive, steps with expensive reagents. We sought to address these challenges using SPOTs’ capabilities in small volume handling, parallel processing, and cross compatibility with existing experimental footprints.

We adapted the protocol of Chen *et al.* for high-throughput library preparation of the 16S V4 region^66^. We first grew colonies on agar plates from environmental samples and suspended one colony in water in each well of a 384 microwell plate. The colonies were heat-lysed and underwent freeze-thaw cycles. The lysate microwell plate was sandwiched to a SPOTs plate, and the liquid was metered to the SPOTs plate by centrifuging it into contact then separating the plates (Fig. 4A, Detailed methods in Supplementary information section S4.1). Polymerase chain reaction (PCR) master mix and indexed primers were loaded onto additional SPOTs plates with row/column loaders (Fig. 1D) to incorporate four indices for multiplexing (Fig. 4, B and C, Supplementary Table II-VI). All reaction components (Supplementary Table VII) were transferred from SPOTs into a second 384 microwell plate by centrifugation, resulting in a total reaction volume of 3 μL/well. Near complete transfer is obtained when centrifuging from SPOTs to microwell plates at sufficient g force. After thermal cycling under oil (to prevent evaporation), the PCR products were pooled by centrifuging into a collection tray spray coated with the same omniphobic coating. The pooled libraries were cleaned and sequenced. Sequences were processed and filtered into representative amplicon sequence variants (ASVs).

**Fig. 4.**
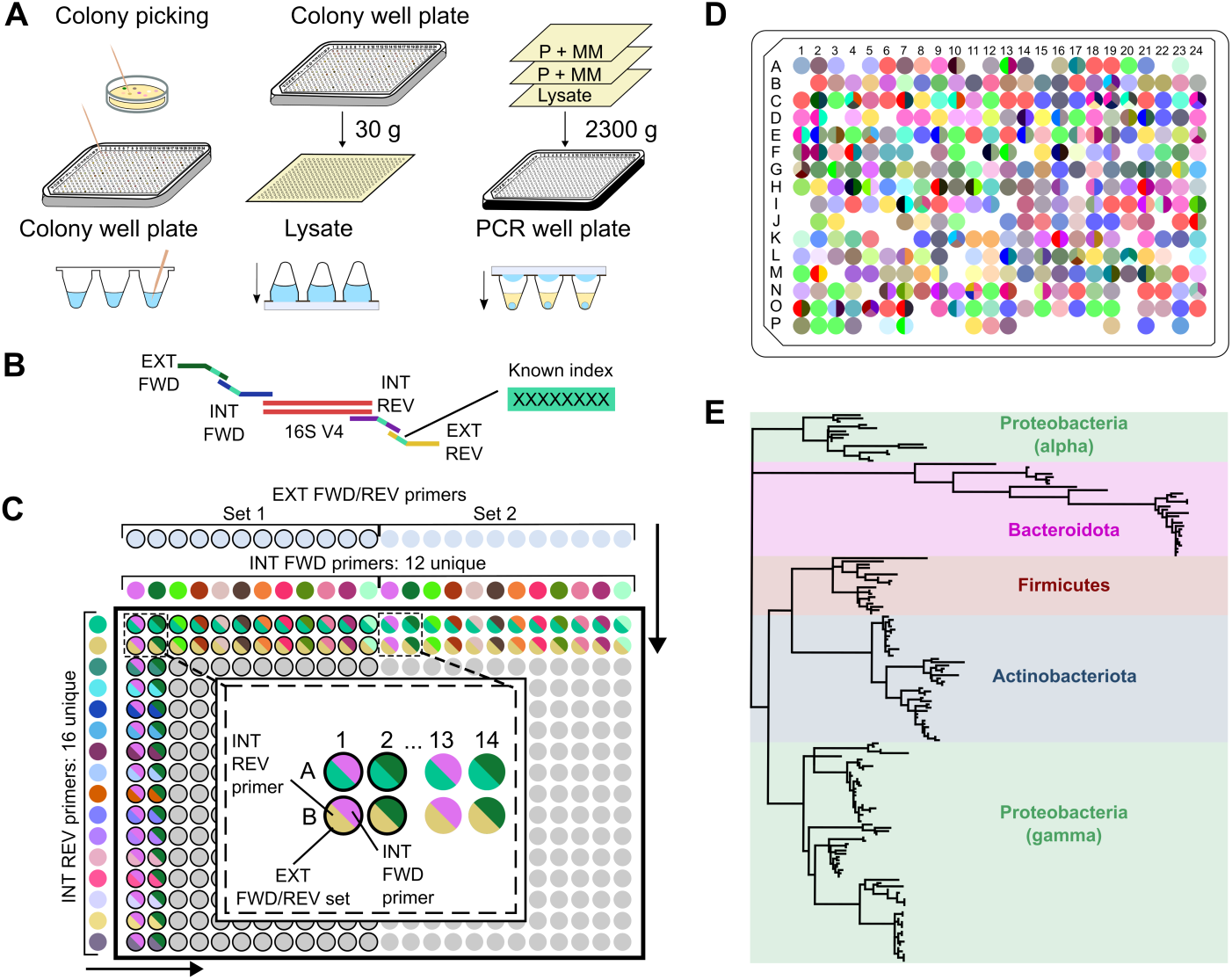
High-throughput microbial genotyping using SPOTs. **a**, Individual colonies were picked from agar plates into a 384 microwell plate and lysed. A SPOTs plate was used to harvest lysate volumes from each well in parallel through centrifugation. Lysate was transferred from the SPOTs plate to a second microwell plate by centrifugation. Mixtures of PCR master mix and primers were loaded on SPOTs plates via row/column loaders and transferred to the second microwell plate by centrifugation. Reactions were thermal cycled under oil to prevent evaporation, then pooled and sequenced. **b,** A four-index primer scheme targets the bacterial 16S rRNA V4 region for amplification and encodes well positions. **c,** The library indexing scheme encodes columns with interior forward primers and rows with interior reverse primers. Two dual-unique exterior primer sets encode the left and right half of the plates. The entire 384 microwell plate is indexed with 32 primers. **d,** Zero to four representative ASVs were found in each well. Each unique color represents a distinct ASV. **e,** A phylogenetic tree shows the 16S taxonomy of sequenced organisms.

We obtained representative ASVs for 88% of wells (338/384), similar to the success rate of 25 µL reaction protocols performed with pipettes^66^. Wells without representative ASVs (12%, 46/384) likely arose from colonies that failed to amplify, which can be due to factors such as fungal colonies (which do not have a 16S rRNA gene), PCR inhibitors, and the colony’s susceptibility to lysis^67,68^. Of the wells with sequences, 77% (260/338) had a single representative ASV and 23% (78/338) had more than one representative ASV (Fig. 4D). The presence of multiple ASVs in some wells may be attributed to mixed colonies or intragenomic 16S variation^66,69^. ASVs recurring in multiple wells are likely attributed to redundant sampling of organisms during colony picking. In total, we obtained 171 representative ASVs from 71 unique genera across four phyla: Proteobacteria, Actinobacteriota, Bacteroidota, and Firmicutes (Fig. 4E), showing how SPOTs can scale this workflow by shrinking volumes and increasing throughput.

More broadly, this final application demonstrates the ability of the SPOTs platform to execute multi-step biological protocols, handle enzymes and DNA, use high-throughput sequencing as a readout, and interface with conventional microwell plates, unlocking a rich landscape of experimental possibilities.

## Discussion

Successful implementation of these three experimental workflows establishes SPOTs as a versatile platform. Our experiments demonstrated different paradigms of experimental design including pairwise combinations, higher order combinations, and parallel processing. We showed how information can be collected from SPOTs experiments using microscopy and sequencing readouts, and how SPOTs can be used as a standalone platform or in conjunction with conventional microwell plates to leverage existing instrumentation. These, and additional modes of use and readouts, can be strung together to facilitate myriad experimental workflows. SPOTs’ profile of technical capabilities, accessibility, and versatility, promise to be enabling for generating large high-quality datasets required to explore complex systems in a wide variety of domains.

## Acknowledgements

We thank R.C. Stokes and H. Vahabi for helpful guidance. Sequencing was performed by the Biotechnology Resource Center (BRC) Genomics Facility (RRID:SCR_021727) at the Cornell Institute of Biotechnology.

