## Supplementary Information for "Surface Patterned Omniphobic Tiles (SPOTs): a versatile platform for scalable liquid handling"

### 1. The SPOTs platform

#### 1.1. Superomniphobic coating

The superomniphobic coating was formulated using a modified procedure from *(26)*. 300 mg of fumed silica particles (0.2 - 0.3 μm, avg. particle size) (Sigma Aldrich) were added to 10 mL of n-Hexane (Sigma Aldrich) and vortexed. The particle suspension was subsequently sonicated in a water bath for 30 minutes and vortexed again to resuspend the settled particles. The resulting mixture turned slightly gelatinous. To functionalize the fumed silica particles, 0.3 mL perfluorodecyl-1H,1H,2H,2H-trichlorosilane (FDTS) (Gelest) was added to the particle suspension. The suspension was vortexed and sonicated again for 30 minutes. Finally, the solution was allowed to equilibrate for three days while occasionally being mixed on a vortex mixer. For spraying, a small amount of the stock coating was diluted in hexane and vortexed. The final diluted suspension was sprayed on a clean glass plate and the loading devices using an airbrush (Paasche) at a pressure of 30 psi held approximately 20 cm from the target surface.

#### 1.2. SPOTs Plates

Clean glass plates were coated with the superomniphobic coating described above. The coating was sprayed until a consistent light haze appeared on the glass. The plates were then baked at 300 °C on a hotplate or in a convection oven for at least eight hours to evaporate any leftover solvent and remove unreacted silane molecules. After baking, the plates were rinsed with deionized water to remove any loose silica particles. The resulting surface was capable of repelling a droplet of up to 70% ethanol-water solution with the surface tension of $\gamma_{lv}$ = 25 mN/m (sometimes the coatings repelled up to 80% ethanol-water solutions). Figure S1 shows that the sliding angles of ethanol solutions of up to 70% (v/v) concentration are very low, making the coatings suitable for the platform in applications involving low surface tension liquids.

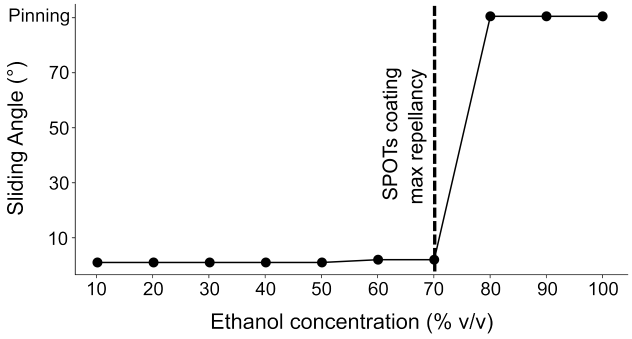


FIG. S1. The sliding angles of aqueous solutions with different ethanol concentrations on a SPOTs plate

To make the desired microdroplet array, predetermined regions of the coated surface were ablated off with an Epilog Fusion Edge 24 laser cutter to create -philic areas that hold desired liquid volumes. The volume of the liquid held by a laser-ablated -philic area is determined by its size and the features of the loading device.

#### 1.3. SPOTs loading devices

Loaders were designed in Autodesk Fusion 360 and machined from Delrin acetal resin sheets (McMaster-Carr) on a Nomad 883 Pro desktop CNC mill (Carbide 3D). Both slot loaders (figure S2a) and row/column loaders (figure S2b) were coated with the superomniphobic coating, described in the coating section, in the same manner as the SPOTs plates. The coated loaders were baked at 120 °C on a hotplate or in a convection oven for at least two hours. The features of the loaders include a slot or hole(s) to hold the liquid, a small step of 0.4 mm to have a consistent gap between the loader and the surface of SPOTs plate, and legs on both sides to keep the loader stably positioned on the plate. Two 1/4" thick stainless steel bars (McMaster-Carr) were clamped onto the top surface of the loader to flatten any bowing in the milled parts. The flattening is necessary to ensure a consistent gap height throughout the loading region, which influences loading volume. After coating, loaders were rinsed to remove any loose particles. 2 mL (slot loader) or 70 μL (row/column loader) of the reagents were pipetted into the slot/hole of the respective loaders to load the plates. To ensure a consistent volume deposition, the row/column loaders should be loaded in the range of 40 μL to 70 μL.

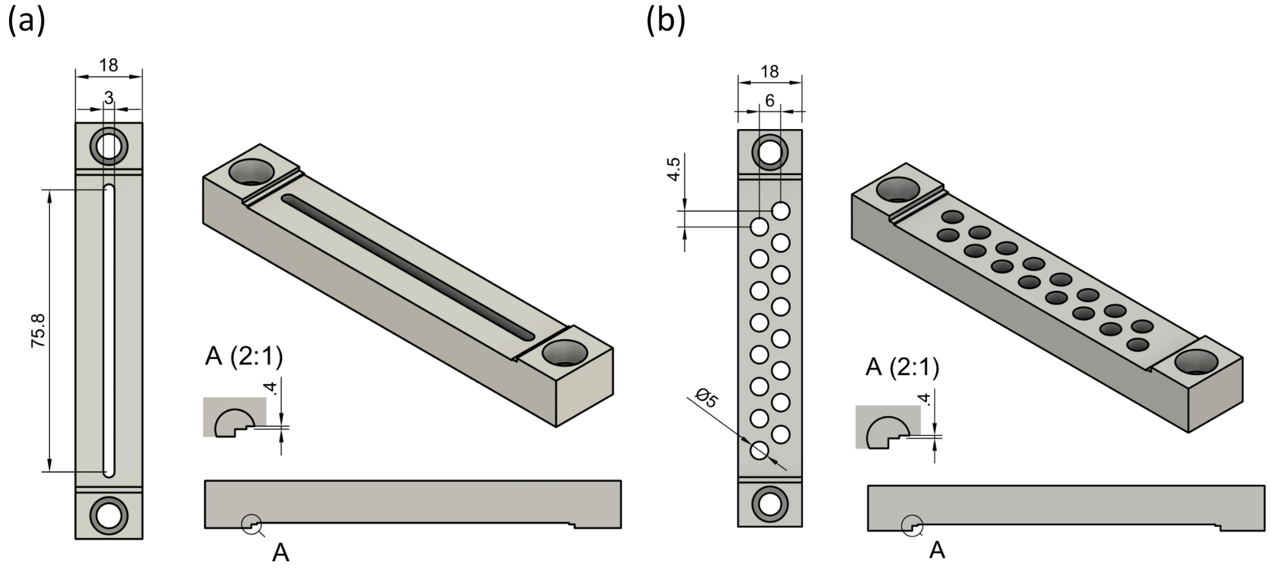


FIG. S2. The critical dimensions of (a) slot loader (b) row/column loader. Loader gap heights are indicated in detailed view A.

#### 1.4. Volume Characterization

Fluorescein disodium salt (Sigma Aldrich) stock solution was prepared at 24 g/L. Fourteen two-fold serial dilutions of the stock solution were performed in a 96-well microwell plate to reach a final volume of 250 μL in each well with four replicates. Subsequently, absorbance readings at 470 nm, 520 nm, and 530 nm were measured using a plate reader (spectro-Max by molecular imaging). Linear regions on the concentration-absorbance curve of each wavelength were used as the calibration curve.

To measure the deposition volume, the fluorescein stock solution was pipetted into the loaders (2 mL in slot loader, 70 μL in row/column loader) and deposited onto the SPOTs plate with -philic areas of known diameters (0.5 mm to 7 mm). A 96 optical microwell plate was pre-loaded with 250 μL of DI water in each well. For diameters 0.5 mm to 4.5 mm, the SPOTs plate loaded with Fluorescein solution was inverted and clamped to the well plate using binder clips. The assembly was then flipped multiple times to ensure complete mixing. For diameters 4.6 mm to 7 mm, the fluorescein solution was pipetted off of the SPOTs plate, and diluted into the microwell plate. The absorbance of each well was measured at 470 nm, 520 nm, and 530 nm. The deposited volume on each -phillic area was calculated using the calibration curve.

##### 1.4.1. Comparison with pipette

Main text figure 1F shows the coefficient of variation (CV) for different volumes on the same plate and different volumes deposited by a single pipette. A second metric of interest is the variation across different plates, and how it compares to the variation across different pipettes. In order to quantify this, we measured 14 volumes across four SPOTs devices (three replicates each) and computed the COV. Similarly, we measured seven volumes (3 replicates each) deposited with three different pipettes. Figure S3 shows these COVs. The COV across different pipettes increases sharply for volumes below 1 µl whereas, across different SPOTs plates, the COV remains below 10% down to 200 nL volumes and below 20% for volumes as low as 9 nL.

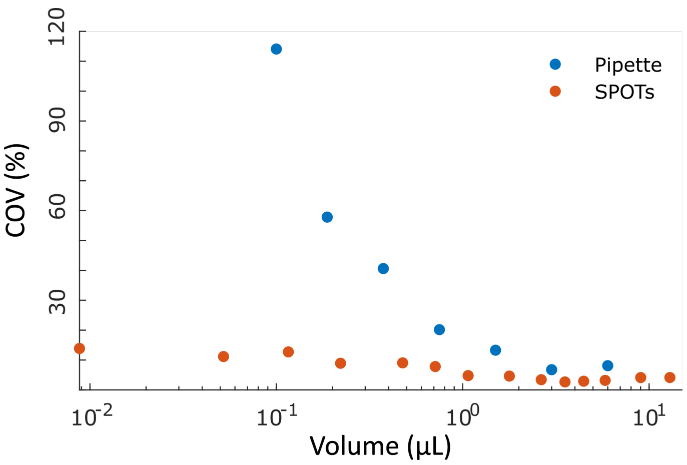


FIG. S3. Global coefficient of variation of volumes deposited by micropipettes (100 nL - 6 µL, COV from 21 volumes across three pipettes, 7 volumes each) and SPOTs platform (9 nL - 13 µL)

##### 1.4.2. Influence of viscosity and surface tension on volume deposition

The influence of physical properties such as viscosity and surface tension of liquids on the deposited volume was evaluated by measuring the deposition volume of glycerol and ethanol solutions of different concentrations. We found that viscosity has minimal effect on the deposition volume (Fig. S4a), and lower surface tension increases deposition volume on smaller -philic areas (Fig. S4b).

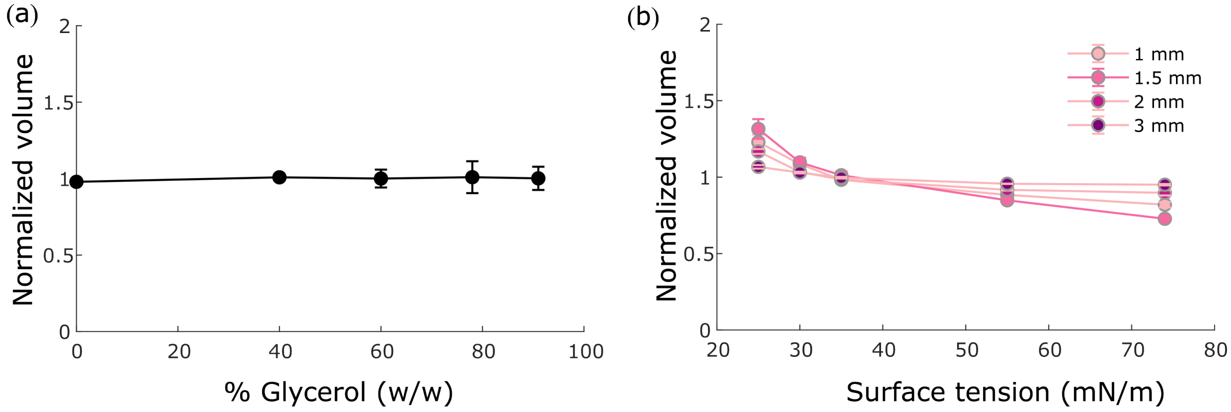


FIG. S4. Influence of physical properties of liquids on volume deposition (a) Volumes of solutions with different viscosities (three replicates of each volume and viscosity) deposited with normal loader sliding speed (∼ 40 mm/s) on a -philic area of diameter 1 mm and (b) Influence of surface tension ($\gamma_{lv}$) on volume deposition on -philic areas of different diameters

The effect of sliding speed on the deposited volumes of liquids with different viscosities was also measured. As seen in figure S5, only a fast sliding speed (200 mm/s) performed on high viscosity liquids has a significant effect on the deposited volume.

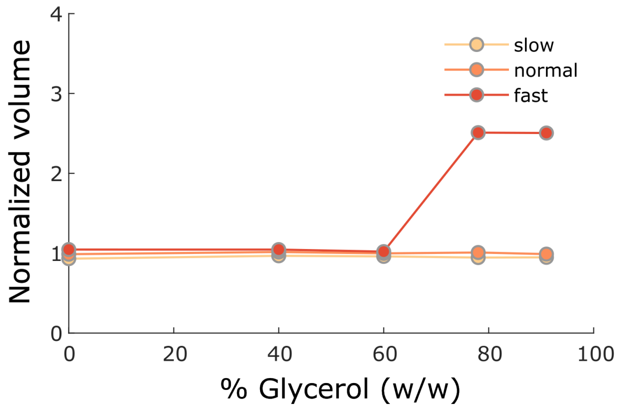


FIG. S5. Influences of sliding speed on the deposition volume of solutions of different viscosities on 1 mm diameter -philic areas (fast: ∼ 200 mm/s, norm: ∼ 40 mm/s, slow: ∼ 10 mm/s) (Note: 'Fast' denotes an excessively high sliding speed, which is impractical for standard experiments)

### 2. Live cell growth and inhibition assays

#### 2.1. Materials and Methods

A single colony of *E. coli* strain W3110 was grown overnight in liquid Luria-Bertani, Miller (BD Difco) (LB) broth. The optical density (OD) 600 reading was obtained with a SpectraMax Plus 384 (Molecular Devices) to dilute the *E. coli* concentration to $1 \times 10^6$ colony forming units (CFU)/mL. Five antibiotics were dissolved in suitable solvents to create fresh antibiotic stock solutions. Amoxicillin (Sigma Aldrich) (AMX) was dissolved in PBS, pH 6.0 and heated for 1 hr at a concentration of 1,000 µg/mL, Doxycycline hyclate (Sigma Aldrich) (DOX) was dissolved in water at a concentration of 5,000 µg/mL, Clarithromycin (Sigma Aldrich) (CLA) was dissolved in acetone at a concentration of 30,000 µg/mL, Trimethoprim (Sigma Aldrich) (TMP) was dissolved in 0.05 M HCl at a concentration of 1,000 µg/mL, and Polymyxin B sulfate (EMD Milipore Corp) (PMB) was dissolved in water at a concentration of 3,000 µg/mL. Each of these concentrated stock solutions were diluted in LB broth to create working stock solutions which were used to perform serial dilutions and determine the maximum dose required to inhibit this *E. coli* strain. These values were used to determine the maximum doses for each antibiotic used in the experiment shown in Table I below:

| Antibiotic | Abbreviation | Maximum Dose (µg/mL) | % of Maximum Dose |
|---|---|---|---|
| amoxicillin | AMX | 2 | 12.5, 37.5, 75, 100, 125, 150, 175, 212.5, 250 |
| polymyxin B sulfate | PMB | 0.45 | 12.5, 37.5, 75, 100, 125, 150, 175, 212.5, 250 |
| doxycycline hyclate | DOX | 0.6 | 12.5, 37.5, 75, 100 |
| clarithromycin | CLA | 36 | 12.5, 37.5, 75, 100 |
| trimethoprim | TMP | 0.3 | 12.5, 37.5, 75, 100 |

Supplementary Table I. Corresponding maximum doses for each antibiotic with the different percentages of maximum doses run within the experiment

A custom MATLAB script was used to randomly distribute single and pairwise combinations of five antibiotics at four concentrations with three replicates across the SPOTs plate. The MATLAB script outputs the design for the receiver plate using 2 mm diameter -philic areas and five different antibiotic plates with a maximum size of 2 mm and a minimum size of 0.5 mm. After the plates were patterned with the laser, LB broth was loaded onto a receiver plate. Subsequently, each antibiotic solution was loaded onto its respective antibiotic plate and sequentially sandwiched onto the receiver plate with a 400 µm gap and mixed by gentle up and down movements. With the receiver plate now consisting of LB broth and all the antibiotics mixtures, a separate plate with the -phillic area pattern matching the receiver plate was loaded with $1 \times 10^6$ CFU/mL of *E. coli* W3110. This plate was then sandwiched together with the receiver plate, separated by a 500 µm gap. The edges of these two plates were sealed with paraffin wax to prevent any evaporation. The plates were then placed in a Nikon Ti2 Eclipse microscope and imaged using dark-field at 35 °C for 21 hours to measure cell growth as an increase in light scattering. Growth for all

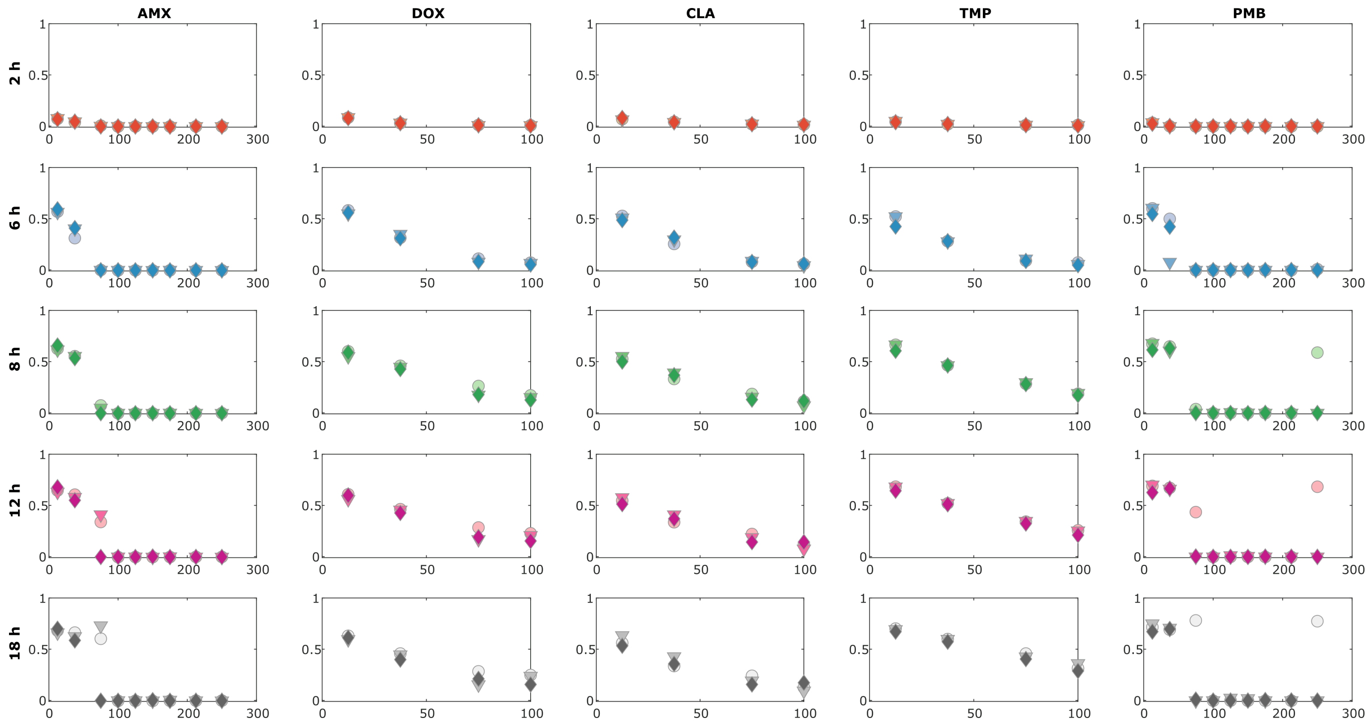


FIG. S6. Single antibiotic growth of scattered light intensity vs concentration for all doses of all antibiotics tested on the experiment plate at time points of 2 h, 6 h, 8 h, 12 h, and 18 h.

doses for each antibiotic tested on the plate are shown in figure S6.

#### 2.2. Analysis

The Bliss scores for pairwise interactions were calculated where: Bliss score $= I_{ab} - I_a \times I_b$. Here, $I_{ab}$ is the intensity of scattered light in the presence of both drugs, $I_a$ is the intensity of scattered light in the presence of drug 'a', and $I_b$ is the intensity of scattered light in the presence of drug 'b'. The bliss scores for all the pairwise combinations of antibiotics at different time intervals are shown in figure S7.

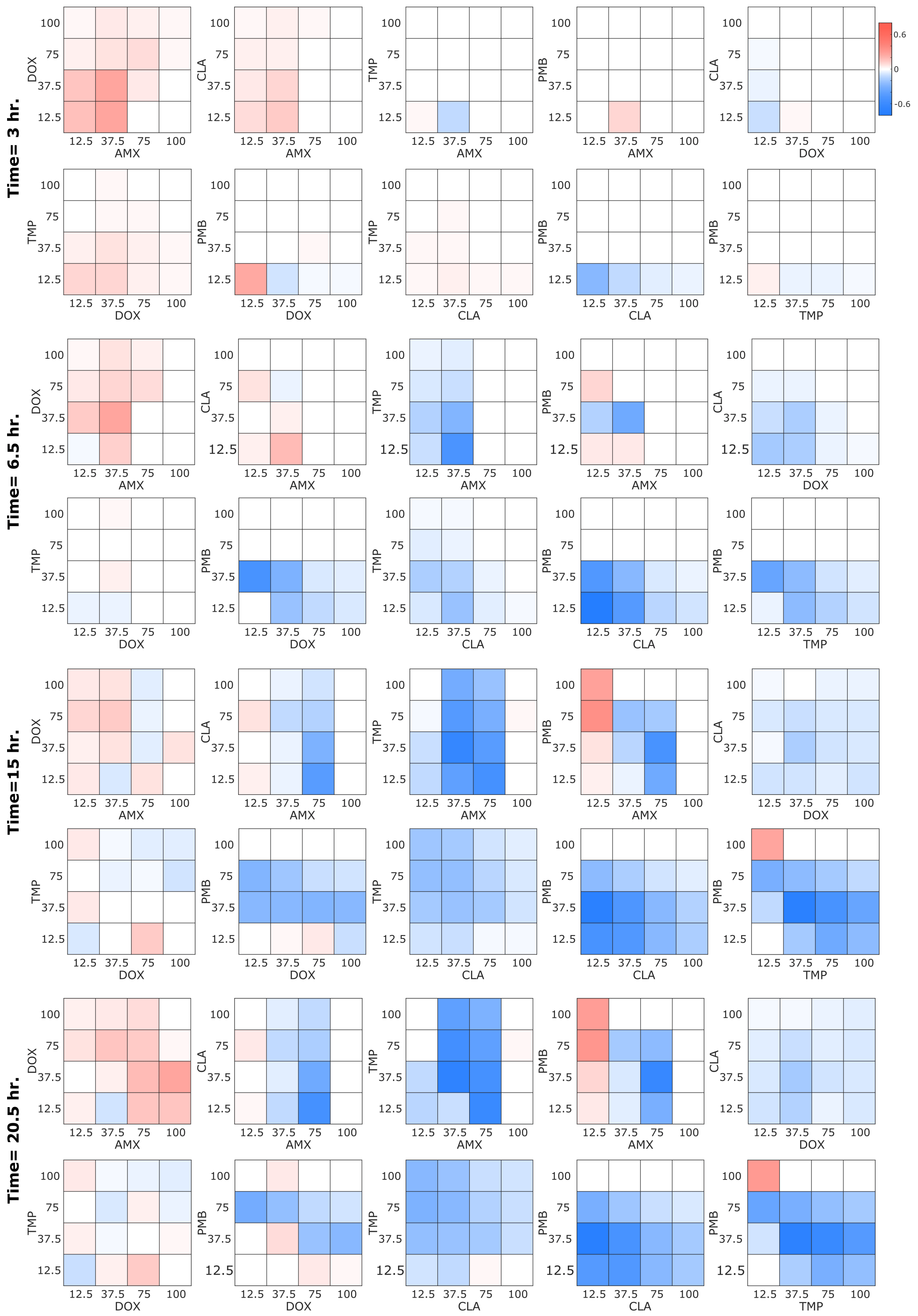

FIG. S7. Bliss scores of concentrations of one antibiotic vs concentrations of another antibiotic for all pairwise combinations of antibiotics at 3 h, 6.5 h, 15 h, and 20.5 h.

### 3. Combinatorial material screening

#### 3.1. Materials and Methods

Stock solutions were prepared for the crystallization of perovskites with the stoichiometric ratio of $ABX_3$. For A site substitutions, five different cations were chosen, for X site substitutions three different anions (halides) were chosen, and for B site, one cation was used. The amounts corresponding to 0.2 M concentration of respective precursors (AX and $BX_2$) were weighed and mixed in dry form and then dissolved together in DMSO. The solutions were allowed to stir overnight. The salts contributing cations and one of the anionic X sites included methylammonium chloride (MACl) (Merck), formamidinium chloride (FACl) (Sigma Aldrich), cesium chloride (CsCl) (Sigma Aldrich), phenethylammonium chloride (PEACl) (Sigma Aldrich), rubidium chloride (RbCl) (Sigma Aldrich) and methyammonium bromide (MABr) (Sigma Aldrich), formamidinium bromide (FABr) (Tokyo chemicals), cesium bromide (CsBr) (Sigma Aldrich), phenethylammonium bromide (PEABr) (Sigma Aldrich), and rubidium bromide (RbBr) (Sigma Aldrich), methyammonium iodide (MAI) (Dyesol) and foramamidinium iodide (FAI) (Dyesol), cesium iodide (CsI) (Sigma Aldrich), phenethylammonium iodide (PEAI) (Sigma Aldrich), and rubidium iodide (RbI) (Sigma Aldrich). For cation B and two of the X site substitutions, the salts included $PbCl_2$, $PbBr_2$, $PbI_2$ all of which were supplied by Sigma-Aldrich. All the salts were used as received from the supplier. Each AX and its corresponding $BX_2$ salts were mixed in a 1:1 molar ratio. For example, MACl was mixed with $PbCl_2$, MABr with $PbBr_2$, MAI with $PbI_2$, and so on, giving fifteen mixtures listed as follows.

1. MACl + $PbCl_2$ ($MAPbCl_3$)
2. MABr + $PbBr_2$ ($MAPbBr_3$)
3. MAI + $PbI_2$ ($MAPbI_3$)
4. FACl + $PbCl_2$ ($FAPbCl_3$)
5. FABr + $PbBr_2$ ($FAPbBr_3$)
6. FAI + $PbI_2$ ($FAPbI_3$)
7. CsCl + $PbCl_2$ ($CsPbCl_3$)
8. CsBr + $PbBr_2$ ($CsPbBr_3$)
9. CsI + $PbI_2$ ($CsPbI_3$)
10. PEACl + $PbCl_2$ ($PEAPbCl_3$)
11. PEABr + $PbBr_2$ ($PEAPbBr_3$)
12. PEAI + $PbI_2$ ($PEAPbI_3$)
13. RbCl + $PbCl_2$ ($RbPbCl_3$)
14. RbBr + $PbBr_2$ ($RbPbBr_3$)
15. RbI + $PbI_2$ ($RbPbI_3$)

In this way, fifteen stock solutions were made, but due to poor solubility of FABr + $PbBr_2$ and CsCl + $PbCl_2$ in DMSO, only the remaining thirteen solutions were loaded onto the SPOTs plates for mixing.
A receiver plate with a specifically designed array of -philic sites of uniform sizes (2 mm diameter) was loaded with DMSO. Subsequently, each of the dissolved AB + $BX_2$ stock solutions was loaded onto a separate specifically patterned SPOTs plate and sandwiched onto the receiver plate sequentially. Upon sandwiching and gentle up and down movements, each plate was left on the receiver plate for 5 minutes to ensure proper mixing. The pattern for each SPOTs plate loading AB+$BX_2$ solution was generated using a custom MATLAB script. For the presented experiment, the patterns on the plates were designed such that the mixtures of cations were created in increments of 25% and the increments of anions was 33%. A total of 612 -philic areas, containing as many mixtures, now acted as distinct reaction sites. We had two reaction sites where due to the unavailability of exact ratios, ratios slightly different than the 25% cation increments and 33% anions increments were made. These combinations correspond to #382 (Cs-69.5%, Rb - 25.6%, and MA, FA, Cs - 1.6% each; Cl - 30.4%, Br - 34.7%, I - 34.7% ) and #457 (FA-69.5%, Rb - 25.6%, and MA, Cs, PEA - 1.6% each; Cl - 34.7%, Br - 30.4%, I - 34.7%) on the supplementary excel sheet. We note that this target composition space includes some mixtures with a higher fraction of Rb content than the expected limit of uptake. Such materials are not expected to form homogeneous perovskites. The receiver plate was left on the microscope stage without any covering and each of the reaction sites was imaged at an interval of 90 minutes for 82 hours to monitor the precipitation dynamics. After the solvent evaporation was complete, a custom-made large area imager equipped with a UV LED emitter (Thorlabs) with an excitation wavelength of 374 nm and a camera with an RGB sensor (Basler #acA3800-14uc) was used to image each reaction site with six different exposure times (50, 100, 500, 800, 1000, 1200 milliseconds). We note that this acquisition setup does not allow us to record UV or IR emissions. In addition, images of each reaction site were also taken at different intervals up to the

period of 172 days to monitor the effect of aging on the precipitated material (Note: during the aging studies, a slight fluctuation in the UV light source was observed over the period.)

#### 3.2. Analysis

To extract information of the material formed on each reaction site, we vectorized the RGB values of each image. First, we normalized the RGB images by dividing each pixel channel value by the sum of the channel values of the pixel. We then constructed RGB histogram bins by splitting the RGB cube into 3D voxel bins. Each side of the cube, i.e. each channel, was assigned 8 bins, resulting in a total of $8^3$ bins. For each image, we assigned to each bin the number of pixels which had an RGB value within the range of the bin. We then flattened the histograms into vectors of length $8^3$. We subsequently applied Uniform Manifold Approximation and Embedding (UMAP), with parameters of 15 neighbors and a minimum distance of 0.5, reducing the vector dimensionality to two, facilitating visualization and exploratory analysis.

In another visualization, we created ternary plots to evaluate the effects of each cation in a ternary cation mixture for a particular anion combination as shown in the figure S8.

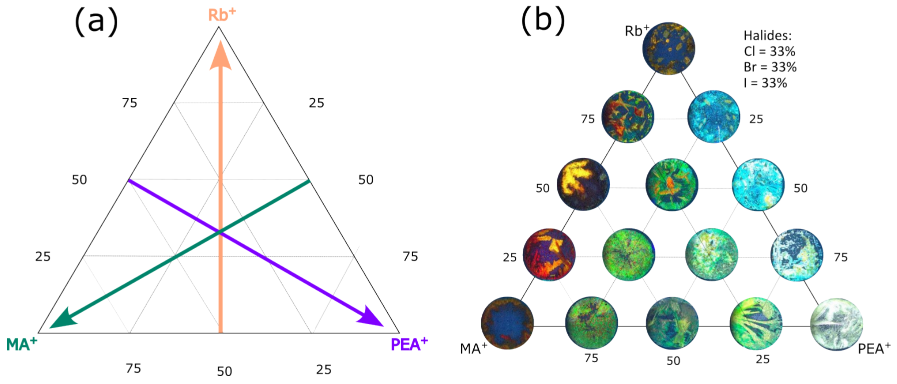


FIG. S8. (a) A schematic ternary plot illustrating the increase of each cation along the respective axis. (b) The ternary plot of $MA^+$, $PEA^+$, and $Rb^+$ ($Cl^-$:$Br^-$:$I^-$ - 1:1:1) showing the variation in the material color as each of the cation fractions increases along the respective axis

For each anion combination, the ternary plots can be combined together to visualize multiple ternary combinations simultaneously, figure S9 shows two such ternary plots combined together.

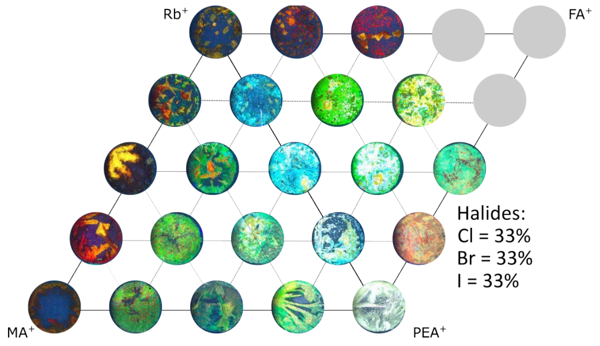


FIG. S9. Two ternary plots combined together with an overlapping axis visualizing the effect of cation fraction variation of $MA^+$, $PEA^+$, $Rb^+$ and $FA^+$ ($Cl^-$:$Br^-$:$I^-$ - 1:1:1) on the material color (Gray empty locations indicate the combinations which could not be made because of the insolubility of two of the mixtures ($FABr+PbBr_2$ and $CsCl+PbCl_2$))

Expanding on this analysis, we optimized the combinations of all the ternary plots formed for ternary mixtures of the five cations used, with maximum overlapping of the axes. This assembly of plots encompasses all the possible ternary cation mixtures for each anion combination. Figure S10 shows the illustration of such assembled ternary plots for 10 anion combinations formed in our experiment. Because of higher emission intensity, images of some of the materials in S10 were saturated, resulting in a white appearance. We created an additional ternary diagram using images captured with lower exposure settings (500 milliseconds), as depicted in S11.

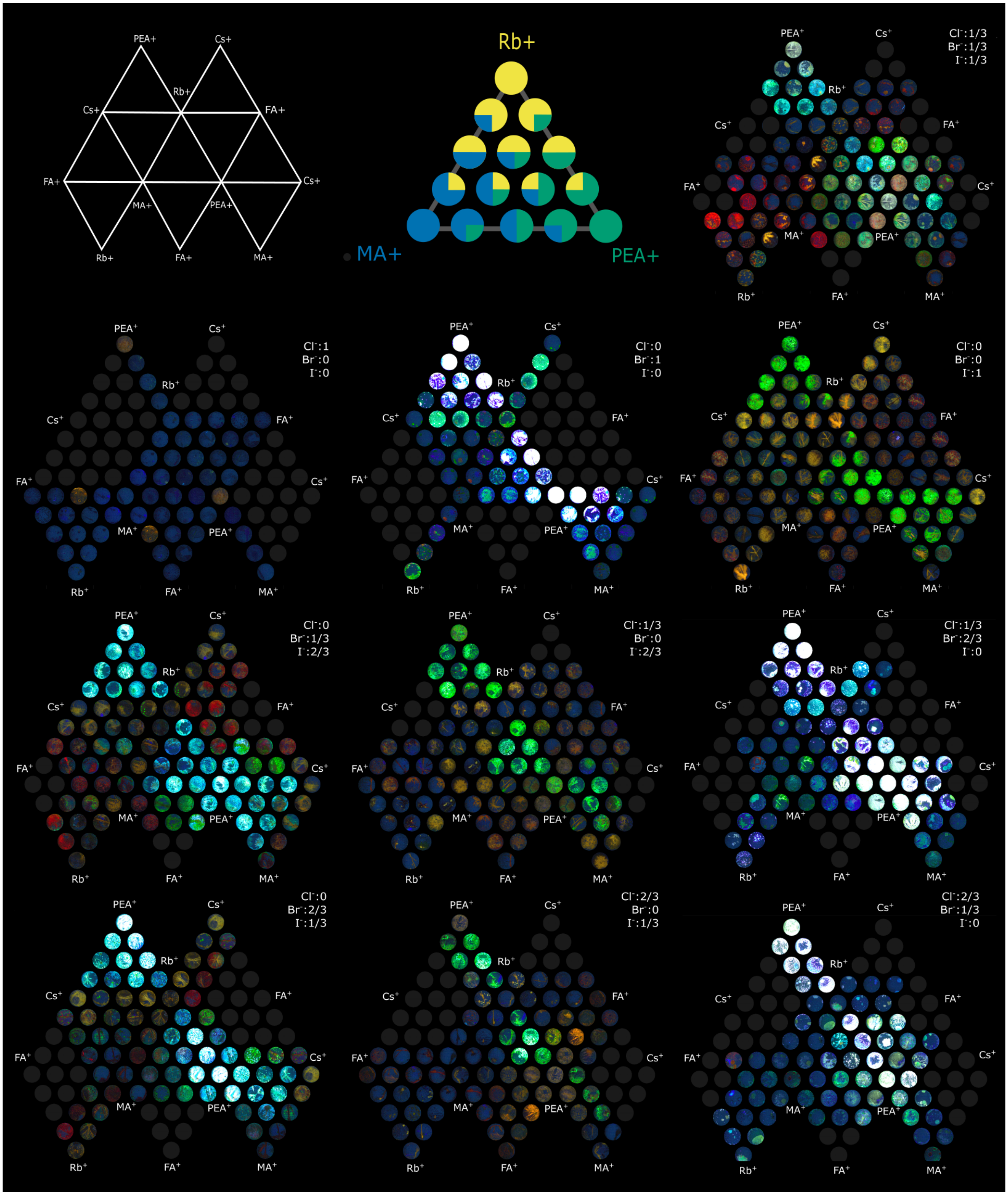


FIG. S10. The combined ternary plots for all of the possible ternary mixtures of the 5 cations for each of the 10 anion ratios (Gray empty locations indicate the combinations which could not be made because of the insolubility of two of the mixtures (FABr+$PbBr_2$ and CsCl+$PbCl_2$)). (Top left - Ternary diagram structure. Top center - Each triangle in the ternary diagram represents materials with three select cations. Cation compositions are varied in increments of 25 %)

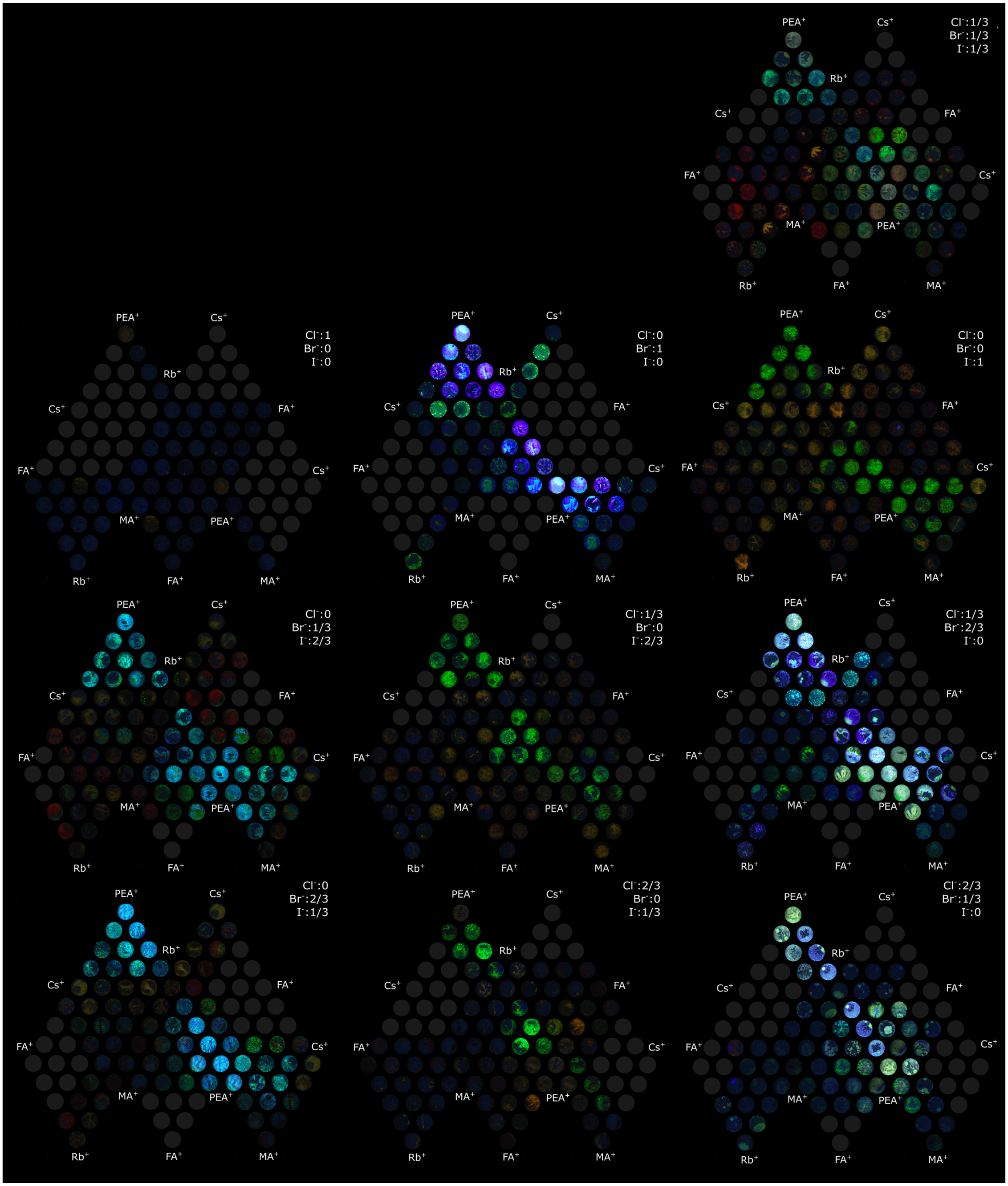


FIG. S11. The combined ternary plots for all of the possible ternary mixtures of the 5 cations for each of the 10 anion ratios (same as S10), with images taken at lower exposure (500 milliseconds) settings.

##### 3.3. Combinations yielding homogeneous colors

For further analysis of compositional trends correlated with each color, we isolated the -philic areas containing materials of homogeneous colors (Red, Blue, and Green) and identified corresponding central compositions and the type and extent of substitutions.

We found that the blue materials predominantly had PEA in the fraction of 0.25 to 1 together with Br. The compositions yielding green materials did not follow any clear trend except that the PEA and I were present in a maximum number of instances. In all the -philic areas where red materials were formed, FA and a combination of Br and I were found to be present. Notably, the presence of Rb up to a fraction of 0.75 was seen to be present in these mixtures without affecting the color homogeneity of the resulting precipitates. A total of 44 locations resulted in green materials, the locations, and the respective compositions which resulted in the homogenous red, blue, and green materials can be seen in S16 and figure S13

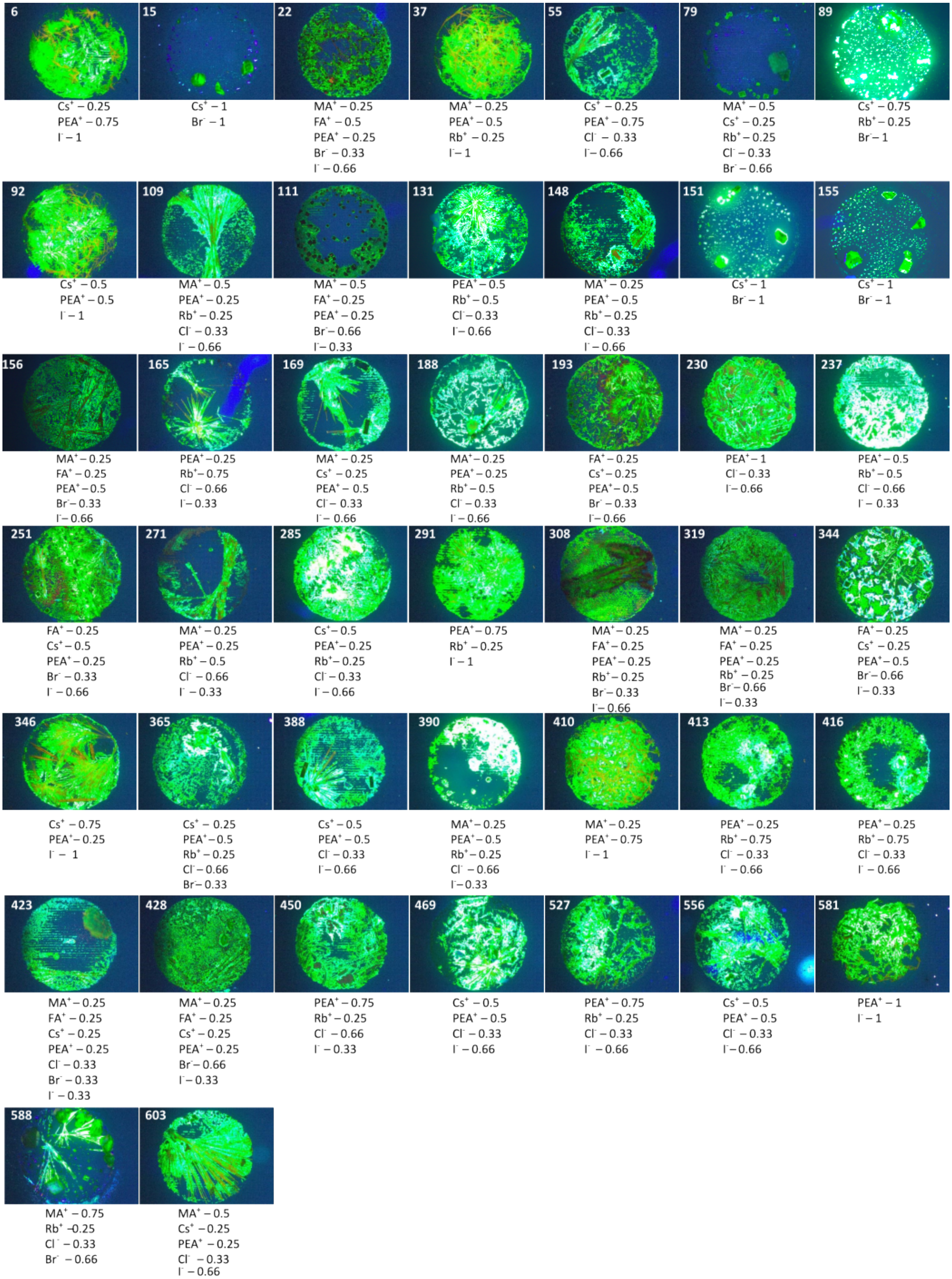


FIG. S12. The set of locations and corresponding compositions which resulted in green material (Note: The images with darker backgrounds are from low exposure settings to visualize the material with high intensity emission)

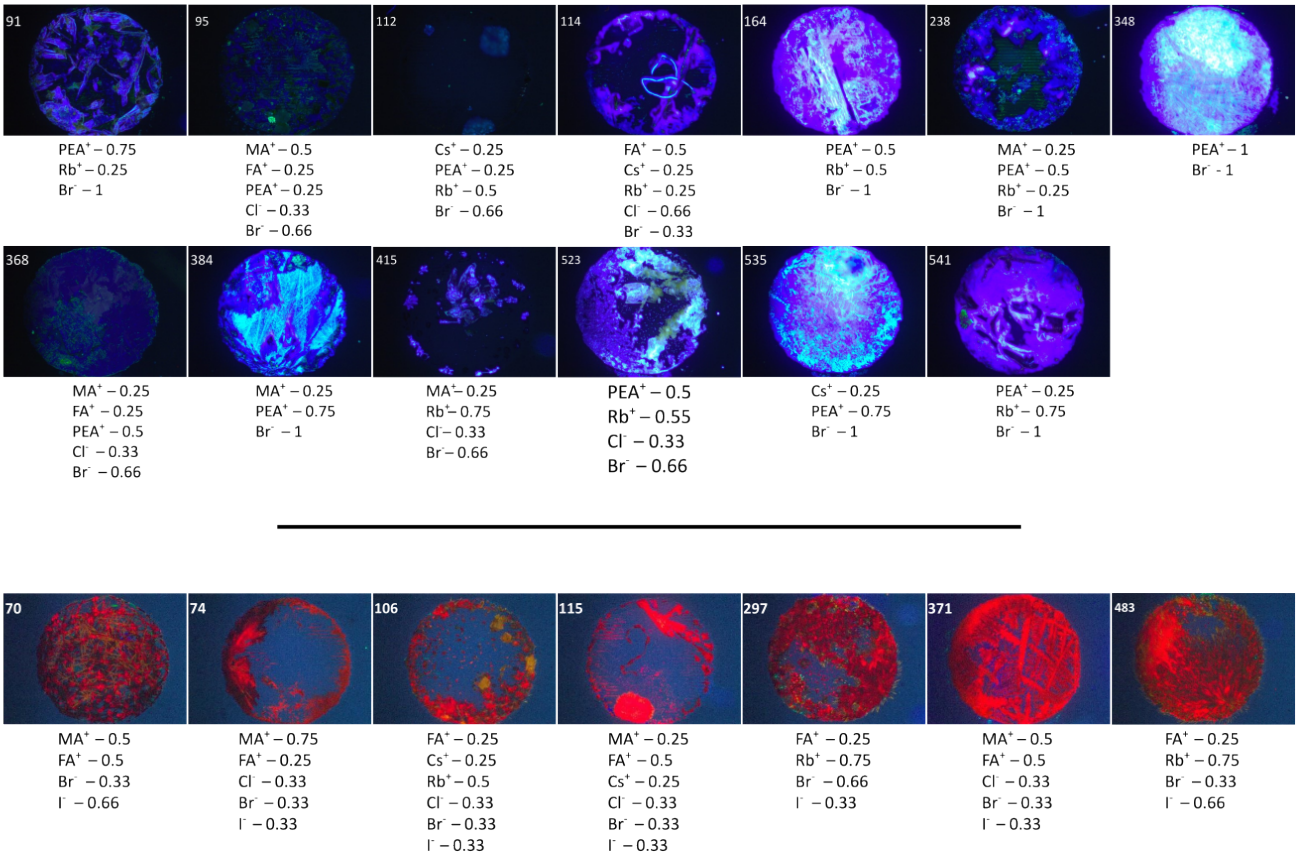


FIG. S13. The set of locations and corresponding compositions which resulted in blue and red material

#### 3.4. Mixing composistions of interest manually

We manually combined a selection of compositions with interesting emission properties in order to compare the outcomes with the mixtures generated on the SPOTs platform. We found that the color of the materials formed after manual mixing corresponds well with the colors formed when mixing is done using SPOTs platform as can be seen in figure S14

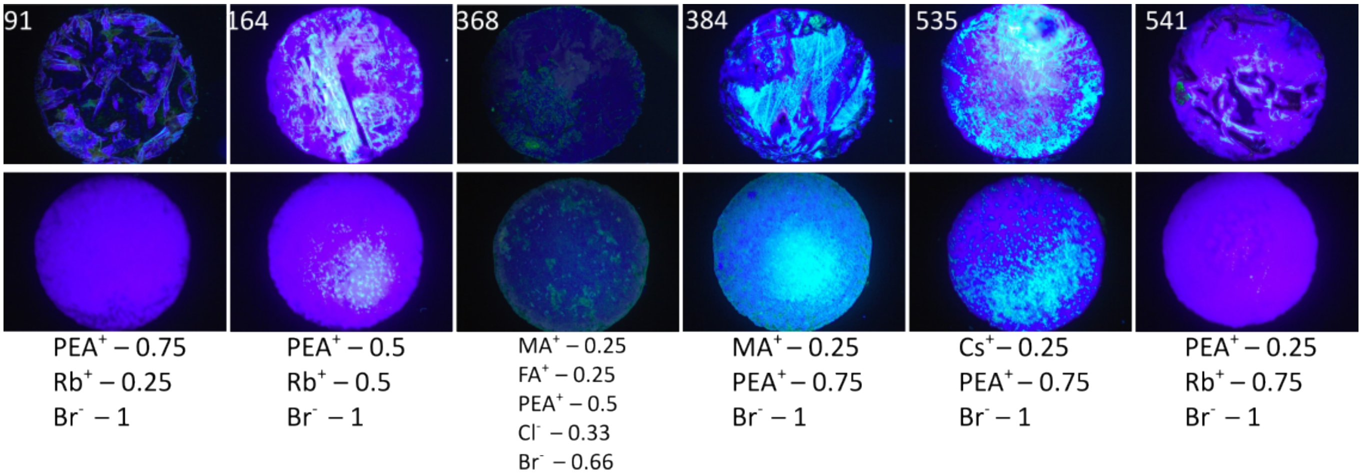


FIG. S14. Example of selected ion compositions mixed on SPOTs plate (top row) and the same composition mixed manually (bottom row)

#### 3.5. Higher resolution sweep for selected compositions of interest

We also made intermediate compositions to see the effect of ion substitutions at a finer resolution. Figure S15 shows the intermediate concentrations containing different ratios of MA, FA, and Cs cations. Here we can see that the increase in the fraction of Cs at the cost of FA in MA, FA, Cs mixtures leads to a loss in intensity of red crystals and on the other hand substitution of MA by Cs in the same ternary cation mixtures does not affect the intensity but the color homogeneity is reduced as the Cs Fraction increases. A similar trend is visible in the photoluminescence measurements performed using 375 nm excitation wavelength and averaging the emission intensity of ten scans (**Note:** The photoluminescence measurements were performed on materials of the same composition but synthesized separately in a different batch). These results demonstrate the application of the platform to locate and narrow down the focus around specific combinations and hence tune the material properties desirably.

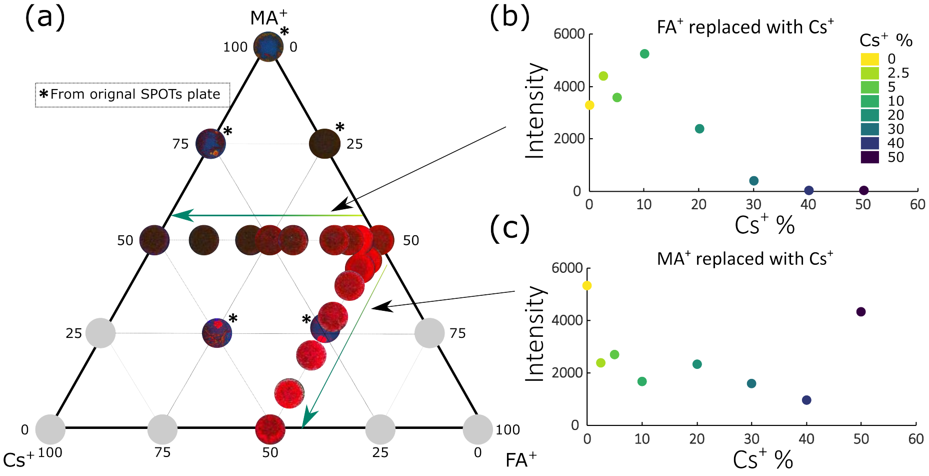


FIG. S15. (a)The ternary mixture of $MA^+$, $Cs^+$ and $FA^+$ ($Cl^-$:$Br^-$:$I^-$ - 1:1:1) with intermediate finer resolution compositions made by doping Cs while replacing MA and FA (b) Intensity versus Cs% for the ternary mixture where FA is replated with Cs (keeping MA constant) (c) Intensity versus Cs% for the ternary mixture where MA is replated with Cs (keeping FA constant) **Note:** Intensities in (b) and (c) taken from the photoluminescence data

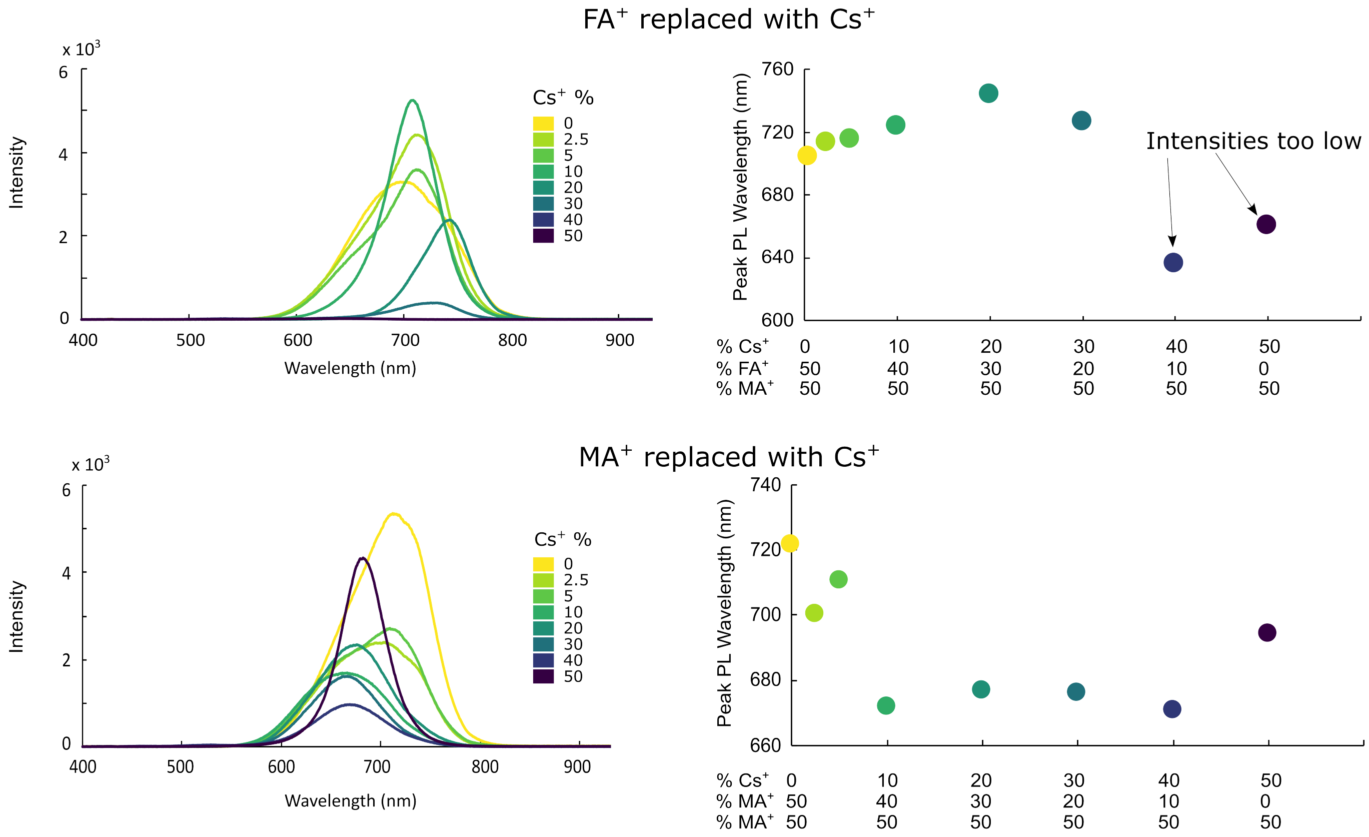


FIG. S16. The photoluminescence spectra of each MA and FA mixtures with gradual doping of Cs ions while replacing FA (top row plots) and MA (bottom row plots)

#### 3.6. Reproducibility within the SPOTs plate

In order to confirm the reproducibility of mixtures formed from the same intended combination at different locations of the plate, we created two additional replicates for 35 of the combinations, and we observe that all the products from the same compositions gave similar results. Figure S17 shows the images of the replicates created and their corresponding compositions.

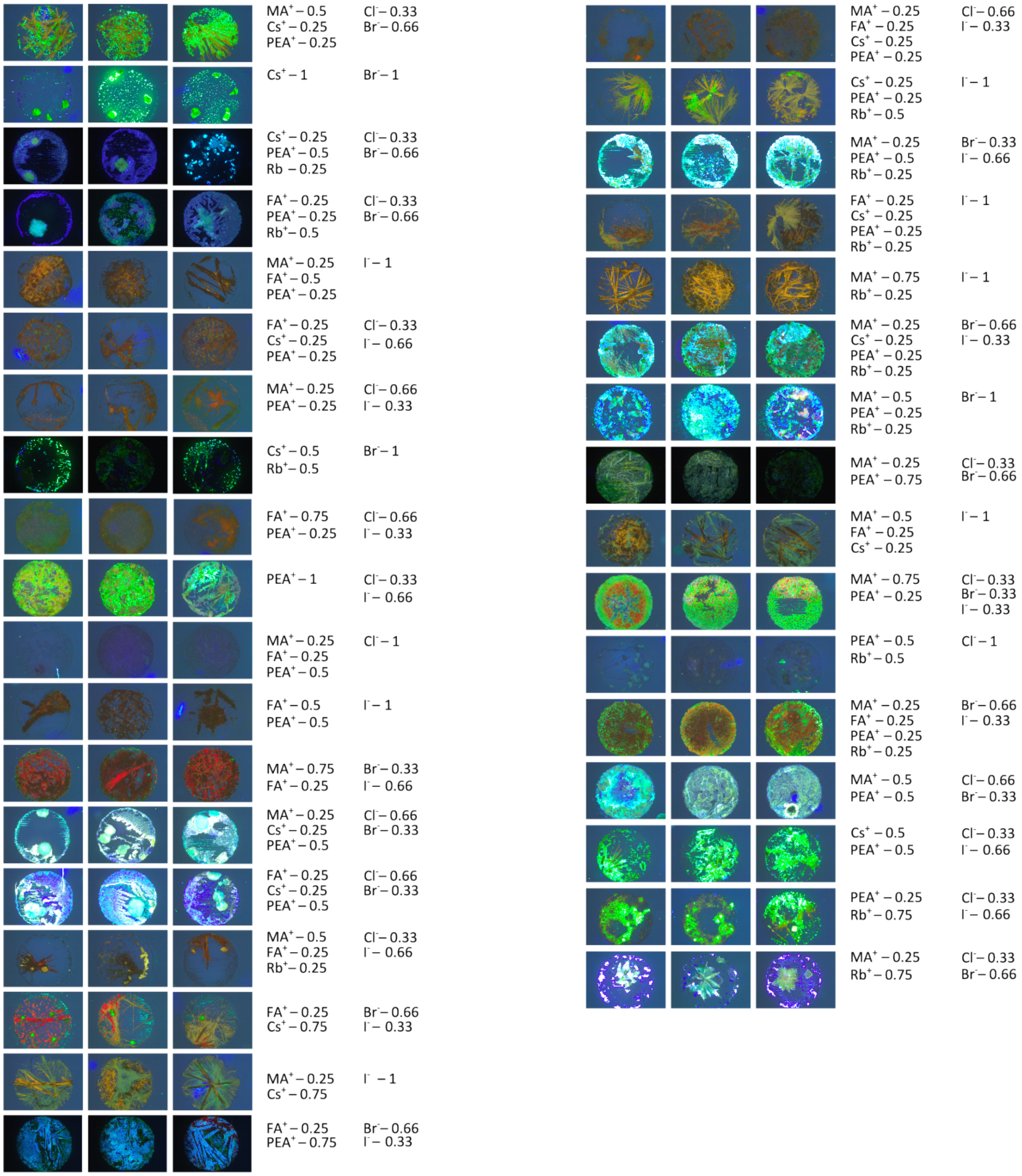


FIG. S17. The images of all the triplicates made on the SPOTs plate and their respective compositions( Note: The images with darker backgrounds are from low exposure settings to visualize the material with high intensity emission)

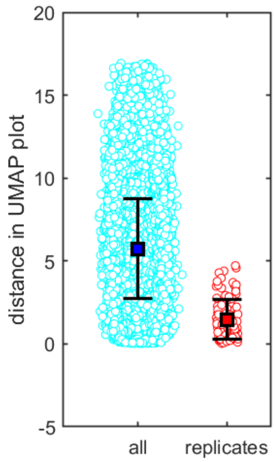


FIG. S18. Distances between materials within UMAP. The replicate materials are on average closer within the UMAP visualization than non-replicate materials

### 4. Genotyping microbial isolates

#### 4.1. Materials and Methods

**Colony preparation:** 20 environmental samples (18 soil samples, 2 creek water samples) were collected from various locations near Cornell University, Ithaca, NY. Each sample was eluted in DI water. Soil samples were diluted by $10^4$ and $10^5$ while water samples were diluted by $10^1$ and $10^2$. 100 µL of each dilution was inoculated on 1x, 0.1x, and 0.01x tryptic soy agar and incubated at 25°C for 7 days. Microbial colonies of different morphology on each agar plate were picked individually with sterile toothpicks. The colonies were suspended in wells containing 20 µL of 0.01% phenol red solution in a 384 microwell PCR plate (VWR). The addition of phenol red allows for visualization of successful liquid transfer to the reaction mix. The microwell plate was heated to 95°C for 5 minutes and subjected to three freeze-thaw cycles at -80°C and 20°C to lyse the microbial cells.

**Primer design:** The PCR primer design was adopted from Chen et al. (Chen et al, 2023) to target 16S V4 rRNA. Briefly, the interior forward/reverse primers encode the column/row position of each well, respectively. Two sets of dual-unique exterior primers encode the left/right half of the 384 microwell plate. Primer sequences and indices are as follows:

| Primer | Sequence (5' to 3') |
|---|---|
| Interior FWD | TCGTCGGCAGCGTCAGATGTGTATAAGAGACAGXXXXXXXXGTGYCAGCMGCCGCGGTAA |
| Interior REV | GTCTCGTGGGCTCGGAGATGTGTATAAGAGACAGXXXXXXXXGGACTACNVGGGTWTCTAAT |
| Exterior FWD (i5) | AATGATACGGCGACCACCGAGATCTACACXXXXXXXXTCGTCGGCAGCGTC |
| Exterior REV (i7) | CAAGCAGAAGACGGCATACGAGATXXXXXXXXGTCTCGTGGGCTCGG |

Supplementary Table II. Interior and exterior forward and reverse primers. *Note: XXXXXXXX index changes based on encoded rows/columns

| Interior FWD index | Column |
|---|---|
| ATTACTCG | 1, 13 |
| TCCGGAGA | 2, 14 |
| CGCTCATT | 3, 15 |
| GAGATTCC | 4, 16 |
| ATTCAGAA | 5, 17 |
| GAATTCGT | 6, 18 |
| CTGAAGCT | 7, 19 |
| TAATGCGC | 8, 20 |
| CGGCTATG | 9, 21 |
| TCTCGCGC | 10, 22 |
| GCTCAGGA | 11, 23 |
| GTAGAGAG | 12, 24 |

Supplementary Table III. Interior forward indexes and corresponding columns

| Interior REV index | Row |
|---|---|
| ATCACGAC | A |
| ACAGTGGT | B |
| CAGATCCA | C |
| ACAAACGG | D |
| ACCCAGCA | E |
| AACCCCTC | F |
| CCCAACCT | G |
| CACCACAC | H |
| GAAACCCA | I |
| TGTGACCA | J |
| TACTACGC | K |
| ATCACACG | L |
| CACCTGTT | M |
| CTTCGACT | N |
| TGCTTCCA | O |
| AGAACGAG | P |

Supplementary Table IV. Interior reverse indexes and corresponding row

| Ext FWD (i5) index | Column |
|---|---|
| ATATGCGC | 1-12 |
| ATGCTAGA | 13-24 |

Supplementary Table V. Exterior forward indexes and corresponding columns

| Ext REV (i7) index | Column |
|---|---|
| ACGATCAG | 1-12 |
| CCTGAGAT | 13-24 |

Supplementary Table VI. Exterior reverse indexes and corresponding columns

**PCR:** Row and column loaders were fabricated with reservoir positions that match to 384-well formatted SPOTs plates. Plates and loaders were subject to UV irradiation for 15 minutes to eliminate potential contaminations. Next, 1% bovine serum albumin (BSA) (Sigma-Aldrich) was loaded onto all SPOTs plates and allowed to dry. We found that BSA blocking is crucial to successful PCR reactions when metering reagents with SPOTs plates. The loading mixtures for row/column loaders were prepared as below:

| Component | Column (uL) | Row (uL) |
|---|---|---|
| Interior FWD (10 $\mu$M) | 0.6 | 0 |
| Interior REV (10 $\mu$M) | 0 | 0.6 |
| Exterior FWD (10 $\mu$M) | 5.4 | 0 |
| Exterior REV (10 $\mu$M) | 5.4 | 0 |
| Phusion HF Master Mix | 30 | 30 |
| PCR water | 8.6 | 19.4 |
| Total | 50 | 50 |

Supplementary Table VII. Loading mixtures in each hole of the row or column loaders

10 µL of heavy mineral oil (Innovating Science) was deposited into each well of a 384 microwell PCR plate to prevent droplet evaporation. Three 384 well-formatted SPOTs plates were prepared with loading volumes of 1.25 µL, 1.25 µL, and 0.525 µL per -philic area, corresponding to row loading mixture, column loading mixture, and microbial crude lysate. The loading solutions were metered and deposited on their respective SPOTs plates through row/column loaders, then transferred to the 384 microwell PCR plate by centrifugation at 2300g with a plate centrigue (1 minute). The crude lysate was metered to the matching SPOTs plate by centrifuging the liquid into contact with the -philic areas via a salad spinner, separating, then transfering to the 384 microwell PCR plate by centrifugation at 2300g. PCR sealing foil was applied to further prevent evaporation, and the plate was heated to 80°C for 2 minutes, followed by brief centrifugation at 800g. This procedure was repeated twice to eliminate bubble formation during thermal cycling, which may lead to droplet loss. Thermal cycling was performed on a BioRad C1000 Touch thermal cycler (Biorad) with the following program:

| Step | Temperature(°C) | Time (sec) | Cycles |
|---|---|---|---|
| Initial denaturation | 98 | 180 | 1 |
| Denaturation | 98 | 10 | |
| Annealing | 50 | 30 | 35 |
| Extension | 72 | 60 | |
| Final Extension | 72 | 600 | 1 |

Supplementary Table VIII. Thermal cycling program for PCR reactions in 384 microwell plate

**Pooling:** The liquid in the 384 microwell plate was pooled by centrifugation at 800g to a Thermo Scientific Omnitray$^{TM}$ single well plate (Thermo Scientific) coated with the omniphobic coating. This enables complete collection of the PCR products. The mixture was collected in microcentrifuge tubes and centrifuged to separate the covering oil and the pooled PCR product.

**Cleanup and sequencing:** 50 µL of the PCR product was extracted and purified with AMPure XP beads (Beckman Coulter) at a 0.7x bead/sample ratio. Sequencing was performed with the Illumina MiSeq Nano kit to obtain 1 million 250 base pair (bp) paired-end reads at Cornell Institute of Biotechnology (Ithaca, NY).

**Sequencing analysis:** Raw sequencing reads were demultiplexed by both exterior and interior indices. Reads were truncated to 150 bp, then filtered, merged, and grouped into amplicon sequence variants (ASVs) through DADA2 using default parameters. ASVs were matched with well location based on the indexing scheme from Fig 4C. ASVs were marked as representative if the ASV read abundance in the well passed a minimum read threshold of 5 reads and a minimum proportion of 0.1 of the total read abundance of that well. The phylogenetic tree was created by aligning the ASVs with MUSCLE then using MEGA11 to construct maximum likelihood trees with 100 bootstrap replicates. Taxonomy was determined by finding the sequence of the closest match in the SILVA database through BLAST.
